# Activation and substrate specificity of the human P4-ATPase ATP8B1

**DOI:** 10.1101/2023.04.12.536557

**Authors:** Thibaud Dieudonné, Felix Kümmerer, Michelle Juknaviciute Laursen, Charlott Stock, Rasmus Kock Flygaard, Syma Khalid, Guillaume Lenoir, Joseph A. Lyons, Kresten Lindorff-Larsen, Poul Nissen

## Abstract

Asymmetric distribution of phospholipids in eukaryotic membranes is essential for maintaining cell integrity, signaling pathways, and vesicular trafficking. P4-ATPases, also known as flippases, participate in creating and maintaining this asymmetry through active transport of phospholipids from the exoplasmic to the cytosolic leaflet. In this study, we present a total of nine cryo-electron microscopy structures at a resolution ranging from 2.4 to 3.1 Å, along with functional and computational studies of the human flippase ATP8B1-CDC50A complex, describing the autophosphorylation steps from ATP, substrate recognition and occlusion, as well as its regulation by phosphoinositides. Our findings show that the P4-ATPase transport site is filled with water upon phosphorylation from ATP. Additionally, we identify two different autoinhibited states, a closed and an outward-open conformation. Furthermore, we identified and characterized the PI(3,4,5)P_3_ binding site of ATP8B1 in an electropositive pocket between transmembrane segments 5, 7, 8, and 10. Our study also highlights the structural basis of ATP8B1 broad specificity for lipids and identifies a new transport substrate for P4-ATPases, phosphatidylinositol (PI). We report the critical role of the sn-2 ester bound of glycerophospholipids in substrate recognition by ATP8B1. These findings provide fundamental insights into ATP8B1 regulation, the catalytic cycle, and substrate recognition in P4-ATPases.

## Introduction

Lipid flippases actively transport lipids from the exoplasmic leaflet to the cytosolic leaflet of cellular membranes thereby creating and maintaining an asymmetric lipid distribution between the two leaflets^1^. Transbilayer lipid asymmetry of the late secretory pathway is critical for eukaryotic cell functions such as membrane homeostasis, migration, cell signaling, coagulation and morphogenesis^2^. Most eukaryotic lipid flippases belong to the P4-ATPases subfamily of the P-type ATPase superfamily (with the sole known exception of ABCA4)^3,4^. The P-type ATPases share a common architecture with three cytosolic domains that are involved in the autocatalyzed ATP hydrolysis: the nucleotide-binding (N) domain binds ATP, the phosphorylation (P) domain contains a conserved aspartate residue, which is phosphorylated during the transport cycle, and the actuator (A) domain contains a conserved glutamate, which is responsible for the subsequent dephosphorylation of the phosphoenzyme intermediate, that enables the protein to return to its initial state^5^ (Figure S1a). These three domains are linked to the membrane domain, which is typically formed by ten transmembrane segments (TM1-10). TM4 contains a conserved motif, which is specific to each P-type subgroup. For P4-ATPases it is PISL and it constitutes part of the transport binding site and forms a hydrophobic gate preventing lipids from diffusing freely through the protein^6,7^ (Figure S1a).

Substrate transport by P-type ATPases is driven by large conformational changes of the transporter switching between so-called E1 and E2 states that allow alternating access to each side of the membrane. These conformational changes are tightly coupled to the phosphorylation and dephosphorylation reactions performed by the cytosolic domains. The different steps of the P-type ATPase catalytic cycle are described by the Post-Albers scheme^8,9^ (Figure S1b), with the transition from the E1 state to E1P state being the phosphorylation reaction, and the transition from the E2P state to E2 state being dephosphorylation. Unique to P4-ATPases, the lipid transport is linked to the dephosphorylation reaction, where substrate lipid recognition occurs in the outward-open E2P state followed by cognate substrate-induced occlusion triggering dephosphorylation^10–12^.

The human genome contains 14 different P4-ATPases transcribed by the ATP8, ATP9, ATP10 and ATP11 genes^3^. ATP8B1 was originally identified as the causative gene for progressive familial intrahepatic cholestasis 1 (PFIC1), a rare but severe liver disease characterized by impaired bile flow^13^. Milder diseases caused by mutations of the ATP8B1 gene include benign recurrent intrahepatic cholestasis of type 1 (BRIC1) and intrahepatic cholestasis of pregnancy (ICP1). The function of ATP8B1 has been associated with the integrity of the canalicular membrane of hepatocytes^14^, but the exact mechanism is still unclear, in part obscured by discrepancies regarding the lipid substrate specificity. Furthermore, genetic variation at a site upstream of the ATP8B1 coding region has been identified in resilience to Alzheimer’s Disease^15^, and loss-of-function mutations of the closely related ATP8B4 represent Alzheimer risk^16^, but in both cases again through unknown mechanisms.

Initially, ATP8B1 was reported as a phosphatidylserine (PS)-specific flippase, but other studies have since proposed specificity to phosphatidylethanolamine (PE), phosphatidylcholine (PC) or cardiolipin (CL)^17–22^. More recently, ATPase activity measurements using the purified ATP8B1-CDC50A and ATP8B1-CDC50B complexes also lead to contrasting conclusions^23,24^. Previously, we reported that PC, PE and PS were able to stimulate the ATPase activity, but not CL or sphingomyelin (SM)^23^, whereas Cheng et al., found that PS, but not PC could stimulate activity^24^. Additionally, ATP8B1 activity is tightly regulated by its N- and C-termini that tightly interact with the cytosolic domains in an autoinhibited E2P state^23,24^. In addition to autoinhibition, ATP8B1 activity depends not only on a substrate lipid, but also on the presence of phosphoinositides, with a strong affinity for the triple-phosphorylated form^23^, PI(3,4,5)P_3_.

Here, we present nine different cryo-EM structures, functional studies, and molecular dynamics simulation data of the ATP8B1-CDC50A complex in conformations covering most of the transport cycle and different substrate complexes. From our data we identify the presence of an unprecedently described water molecule associated with the E1 states of the cycle and located in the canonical transport site of P-type ATPase. We also report that the autoinhibited ATP8B1-CDC50A complex oscillates between an outward-open and a closed conformation, investigate a PI(3,4,5)P_3_ binding site, identify new substrates and reveal structural determinants of a broad glycerophospholipid specificity of ATP8B1.

## Results

### Cryo-EM structures of the ATP8B1-CDC50A complex along its catalytic cycle

We expressed the human ATP8B1-CDC50A complex in *Saccharomyces cerevisiae*. Isolated membranes were solubilized with n-Dodecyl β-D-maltoside (DDM) and purified by affinity purification via a BAD-tag (biotin acceptor domain)^25^. DDM was exchanged for Lauryl Maltose Neopentyl Glycol (LMNG) prior to further purification by size-exclusion chromatography. We used two different constructs for our studies: full-length ATP8B1 (WT-ATP8B1) and a variant containing an engineered 3C protease site allowing autoinhibition of ATP8B1 to be relieved by proteolytic cleavage of the C-terminal end (ΔCter-ATP8B1) (Figure S2). The ΔCter-ATP8B1 construct was incubated with nucleotides (AMPPCP or ADP) and/or AlF_4_^-^ to stabilize forms similar to E1-ATP, E1P-ADP and E1P conformations, corresponding to autophosphorylation from ATP and subsequent ADP release events (Figure 1 & S3, table S1). Furthermore, the phosphate mimics BeF_3_^-^ and vanadate were used in presence of transport substrate to stabilize the E2P outward-open and the E2┅Pi lipid-occluded conformation of the complex. Finally, the full-length ATP8B1 preincubated with ATP revealed two off-cycle autoinhibited states.

**Figure 1.**
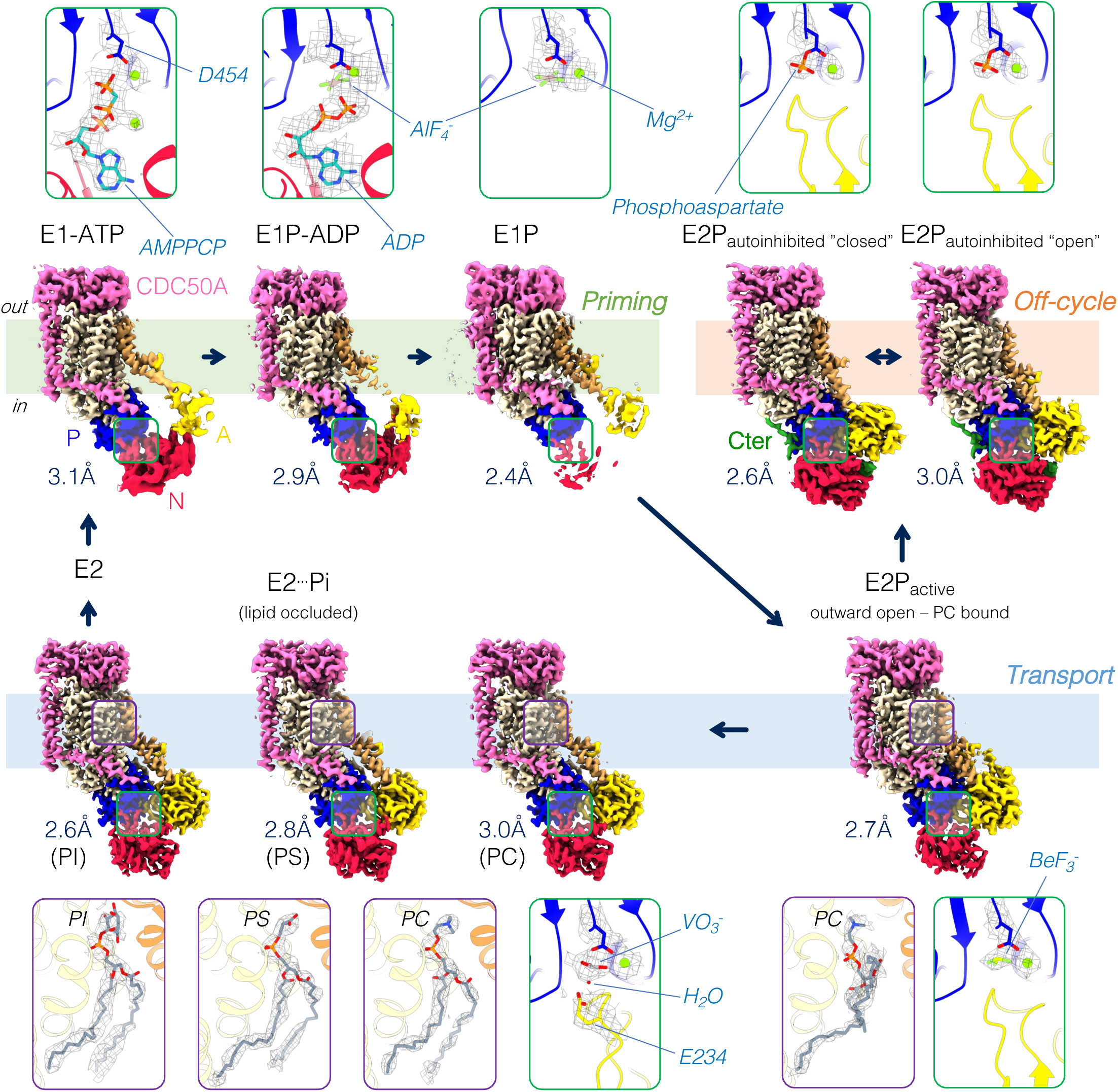
Cryo-EM maps of the ATP8B1-CDC50A flippase complex. Cryo-EM maps of ATP8B1 in complex with CDC50A in different conformations: E1-ATP, E1P-ADP, E1P, E2P_autoinhibited_ “closed”, E2P_autoinhibited_ “open”, E2P_active_, E2┅Pi (PC), E2┅Pi (PS) and E2┅Pi (PI). For each conformation, a close-up view of the phosphorylation site (green frame) and/or of the lipid transport site (purple frame) including their associated EM maps (black mesh) are shown. Color code: The cytosolic A-, N-, and P-domains of ATP8B1 are colored in yellow, red and blue, respectively. The transmembrane domain of ATP8B1 is colored in wheat, with TM1and TM2 in brown. The C-terminal tail of ATP8B1 is colored in green. CDC50A is colored in pink.

### A canonical ion binding site of P-type ATPases is occupied by water in the E1 half-cycle of ATP8B1

The cryo-EM structure of the ATP8B1-CDC50A complex in presence of AMPPCP mimics an ATP bound E1 state and reveals the ATP binding pocket of ATP8B1 (Figure 2a-b). As to be expected, ATP is bound similarly to other P-type ATPases. The adenosine base is stacked by a π-π-interaction with F596. Additionally, D554 is directly engaged in the coordination network of the ß-phosphate, mediated by R652. Indeed, the R652 rotamer state is constrained by the two negatively charged aspartates D554 and D734 (Figure 2b). Noteworthy, the D554N mutation is one of the PFIC1 mutations. The E1P-ADP state mimicked by ADP and AlF_4_^-^ is similar to previously published structures of P4-ATPases^7,12,26^. However, closer inspection of all E1 states along with the outward-open E2P_active_ state reveals the presence of an extra density at the canonical transport site of P-type ATPases between TM4, TM5 and TM6 (Figure 2c). Due to the geometry of the site with no more than four ligands, we model the site as a water molecule. Interestingly, the water molecule is in interacting distance of the S401 from the PISL motif of TM4, the backbone carbonyl group of two residues from TM6 (N989 and T993) and the highly conserved K957 of TM5, which is already described as essential in ATP11C^26^. Given the high conservation of these residues among all P4-ATPases, we compared the cryo-EM densities of other P4-ATPases obtained at a sufficient resolution to observe a coordinated water molecule (Figure S4). In almost all maps, a density could be observed at this site, consistent with a water molecule. The water molecule site partially overlaps with heavy metals and cation sites of P1 and P2-ATPases (Figure S5). Furthermore, the recent cryo-EM structure of human ATP13A2 (a P5-ATPase), in the E1-ATP conformation also revealed water molecules in the canonical P-type ATPase binding site^27^ (Figure S5).

**Figure 2.**
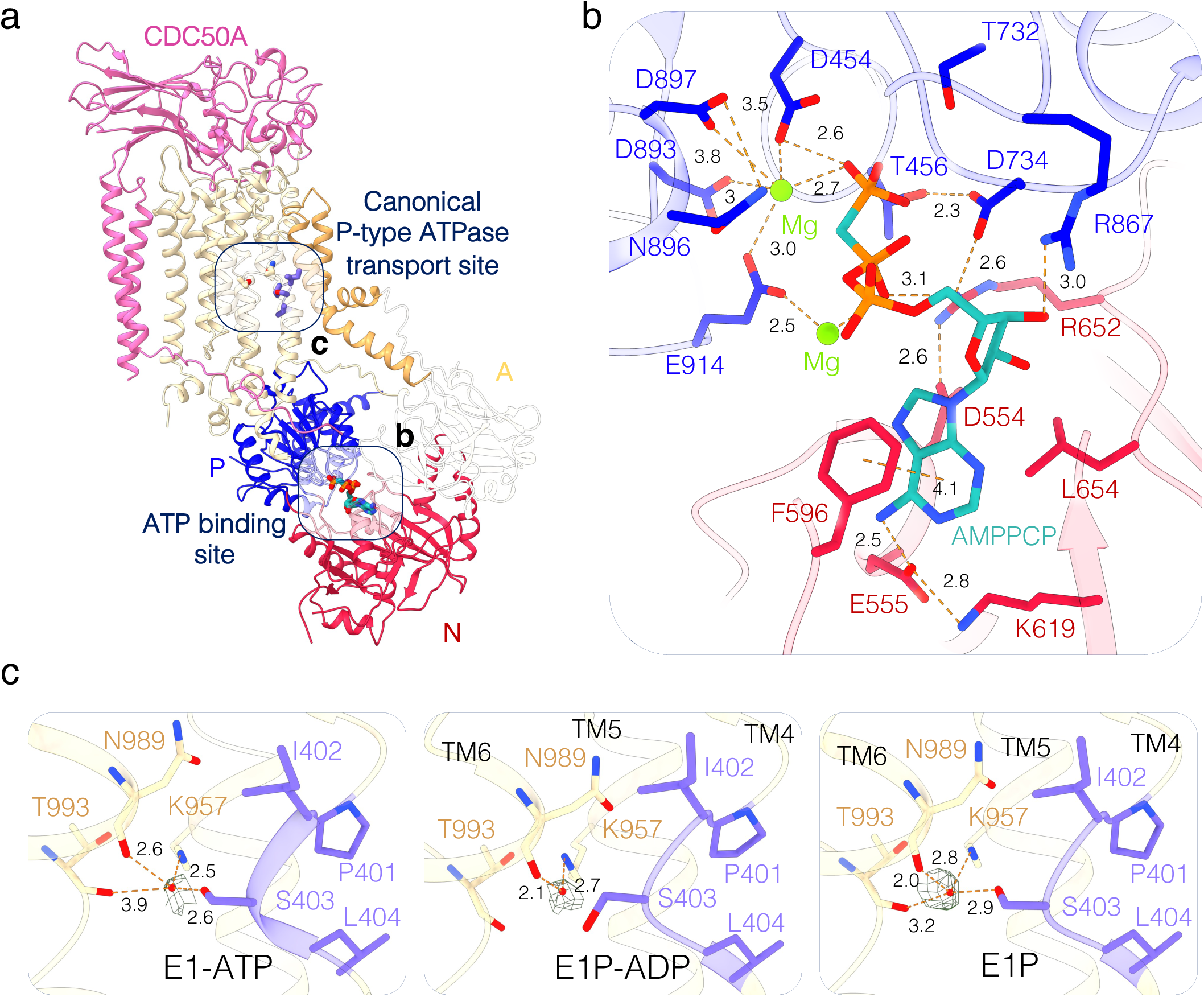
ATP binding and the presence of a water molecule in the core of the lipid transport site. **(a)** ATP8B1-CDC50A structure in a E1-ATP conformation with bound AMPPCP. Color code as Figure 1. **(b)** Close-up view of the AMPPCP binding site including two Mg^2+^ ion with coordinating residues as sticks. **(c)** Detailed views of a part of the not yet formed lipid transport site in the E1-ATP (left, contour map: 4.5), E1P-ADP (middle, contour map: 9.11) and E1P (right, contour map: 7.27) conformations, all revealing the presence of a water molecule (carving distance 1.1Å, for the full EM maps of the region see Figure S4) The water molecule associated EM map is shown in mesh and the P4-ATPase specific PISL motif of TM4 is shown in purple.

### Autoinhibited ATP8B1-CDC50A can adopt an open and a closed conformation

Previously, two independent structures of the ATP8B1-CDC50A complex in the E2P_autoinhibited_ state were reported^23,24^. Surprisingly, they are not identical, especially in the N-terminal part of the transmembrane region (Figure S6a). Here, we collected cryo-EM data of the full-length ATP8B1-CDC50A complex preincubated with ATP and the activating lipid PI(3,4,5)P_3_. Upon data processing, two different conformations were observed (Figure 3). The first one, representing approx. 90 % of the particles, was in a “closed” conformation virtually identical to previous studies of autoinhibited forms^7,23,28^ (Figure 1, 3 and S6a). In this conformation, TM1 and TM2 are tightly packed to the rest of the transmembrane helix bundle, thus masking the substrate recognition PISL motif of TM4 and closing the exoplasmic lipid entry (Figure 3 and S7). However, a significant fraction of the particles adopted an “open” conformation corresponding to the structure reported by Cheng *et al.,*^24^. Here, TM1-2 are rotated and the lipid-binding groove is open as observed for non-autoinhibited P4-ATPases^26,28,29^ (Figure S7). In both cases, we still observe clear densities for the N- and C-terminal tails, interacting with the A, P and N domains in a similar manner (Figure 3). To better understand whether the TM1-2 orientation of this E2P_autoinihibited_ “open” state is different from the active form of ATP8B1 we collected cryo-EM data of the C-terminally truncated ATP8B1 trapped in an E2P conformation with BeF_3_^-^ in presence of a POPC lipid substrate (Figure 1 and 3). In the resulting E2P_active_ conformation, the transmembrane bundle displays the same organization as the E2P_autoinhibited_ “open” conformation with the lipid groove being open and filled with PC (Figure 3). However, in this case, no density associated with the N-terminal tail could be observed in the cryo-EM map. This suggests that the N-terminal tail only interacts with the core protein in the presence of the autoinhibitory C-terminal tail. Given the previously described strong synergistic effect of the N-terminus on C-terminal tail inhibition^23^, our structural data suggest that the N-terminal tail stabilizes the C-terminal tail in the bound form, thus reinforcing the autoinhibition.

**Figure 3.**
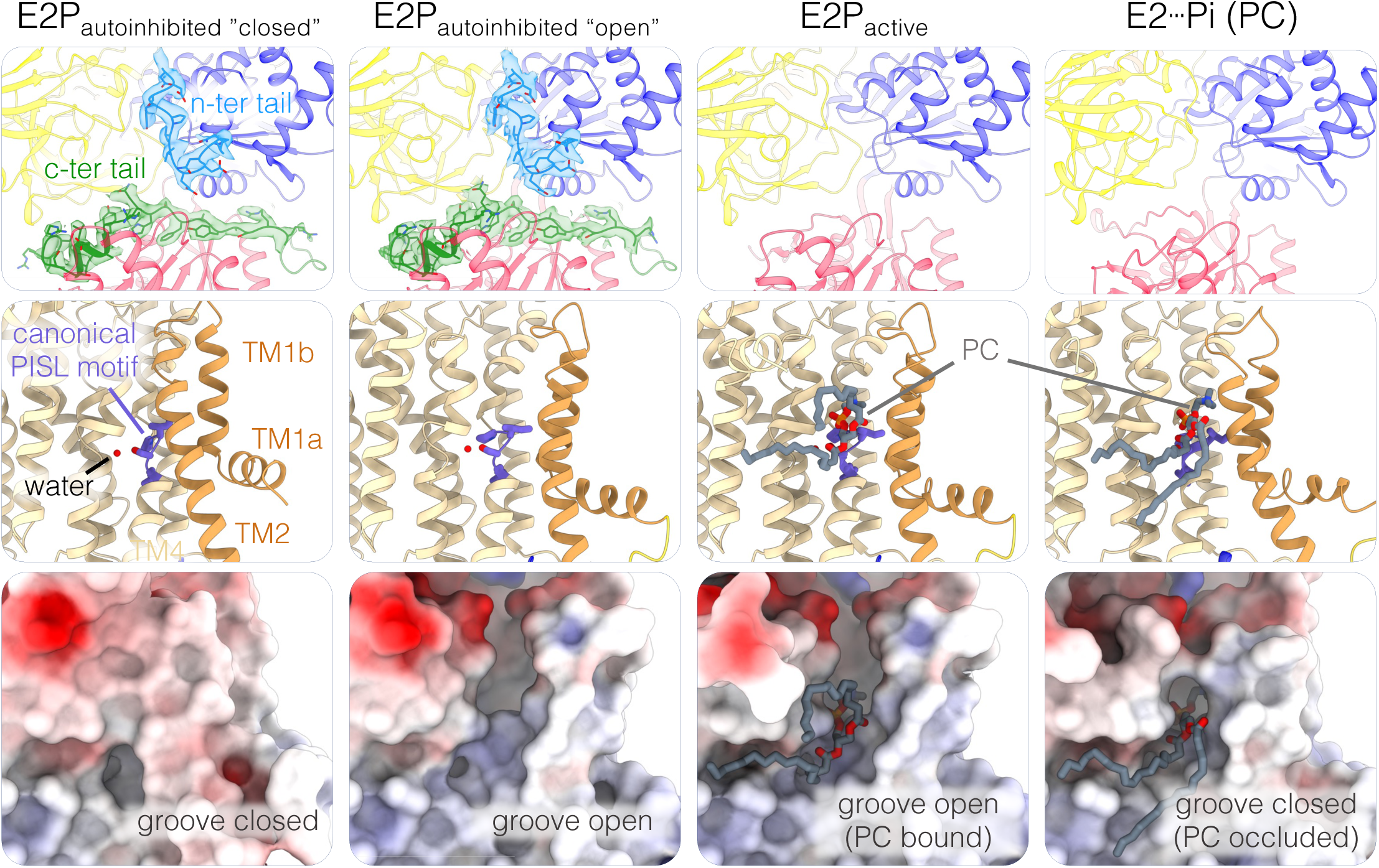
ATP8B1 autoinhibition mechanism and lipid groove opening. Close up view of the cytosolic domains with the ATP8B1 N- and C-terminal tails shown with their associated cryo-EM density when present (top row, contour map: 4.17 and 5.47 for the E2P_autohinhibited_ closed and open conformations respectively) and the lipid binding site in cartoon representation (middle row) or as electrostatic surface (bottom row) in the different E2 conformations. Color code as Figure 1, with the N-terminal tail of ATP8B1 colored in blue.

### PI(3,4,5)P_3_ binding site

As previously showed, C-terminally truncated ATP8B1 exhibited strong stimulation by PI(3,4,5)P_3_ in the presence of a lipid substrate (PC)^23^ (Figure 4a). The cryo-EM structures of the two full-length E2P_autoinibited_ states and the E2P_active_ state were obtained in presence of PI(3,4,5)P_3_ to investigate the mechanism of activation. For the three structures, a lipid density could be observed in a positively charged cavity formed by TM5, TM7, TM8, and TM10 (Figure 4b-c, S8a), similar to the PI(4)P binding pocket of Drs2p^12,28^. The observed density is slightly weaker compared to the surrounding protein, potentially indicating heterogenous occupation of the phosphoinositide binding site and/or flexibility of the ligand. Therefore, we performed molecular dynamics (MD) simulations of two systems based on the structure of the E2P_autoinhibited_ “closed” state: with PI(3,4,5)P_3_ and with a non-phosphorylated PI molecule, matching the position of the modelled PI(3,4,5)P_3_ molecule as a starting point. Both systems contained a native-like lipid composition mimicking a plasma membrane. We then ran a set of five independent 600-ns-long simulations for each system and analyzed the dynamics and interactions between the lipids and the surrounding protein residues. Both the PI(3,4,5)P_3_ and PI remained bound during the entire 600 ns in all simulations, indicating stable binding of the two lipids within the binding pocket. However, during the equilibration procedure, both lipids quickly moved away from the initial positions indicated by the EM densities (Figure S8b). To analyze the interaction between the protein and the lipids, we calculated hydrogen-bond populations over the last 200 ns of each trajectory. Although the number of interacting protein residues is similar for the two systems (12 and 15 for PI(3,4,5)P_3_ and PI, respectively), we found that PI(3,4,5)P_3_ forms more stable contact with the protein compared to PI, as indicated by an overall increase of a hydrogen-bonded population (Figure 4d). For PI(3,4,5)P_3_, the most frequent contacts are established with residues R1032 (0.97), R1164 (0.96) and R952 (0.88). The same residues are also the most frequent interaction partners for PI but with substantially lower populations: R1032 (0.59), R1164 (0.19) and R952 (0.59). This supports a model for the lipid-protein interactions at the PI(3,4,5)P_3_ binding site based primarily on the interactions between positively charged arginine and lysine residues and the negatively charged phosphates on the lipid headgroups. Next, we studied the dynamics of the lipid binding in the cavity by monitoring the positions of the center of mass (COM) of the inositol ring relative to the TM region surrounding the lipid binding site. Indeed, we find that the hydrogen-bonding patterns are closely related to the dynamics of the headgroups belonging to the two different lipids (Figure 4 e-h): PI shows substantially higher mobility in all three directions, whereas PI(3,4,5)P_3_ moves mainly along the z-axis inside the cavity. To visualize this, we clustered^30^ the different binding poses based on the orientations of the inositol. For PI, we find a total of 43 clusters with 19%, 13% and 13% of the structures populating the top three clusters (45% combined), reflecting the large mobility of PI (Figure 4g-h, Figure S8c). In comparison, we find only a total of 9 clusters for PI(3,4,5)P_3_ with 56%, 26%, and 10% of the structures in the top three clusters (92% combined), which is in line with a more stable binding of PI(3,4,5)P_3_. When we examined the solvation properties of the cavity with respect to the bound lipid, we observed on average more water molecules inside the cavity with PI(3,4,5)P_3_ bound (57±9) than with PI bound (47±7) (Figure S8d). Additionally, our simulations indicate that on average 2.7±1.0 K^+^ ions populate the binding site when a PI(3,4,5)P_3_ is present, whereas there are no K^+^ ions are observed in the presence of PI (Figure S8e). This suggests that the PI(3,4,5)P_3_ head group binds additional counterions, despite numerous positively charged residues at the binding site. Altogether, our simulations suggest that the PI(3,4,5)P_3_ binds in a more stable fashion, where dynamics are restricted to mostly “up and down” movements (z-axis), whereas PI exhibits increased 3D mobility. Our MD studies are consistent with the EM maps, which show that PI(3,4,5)P_3_ is the most well-ordered ligand, but is still dynamic.

**Figure 4.**
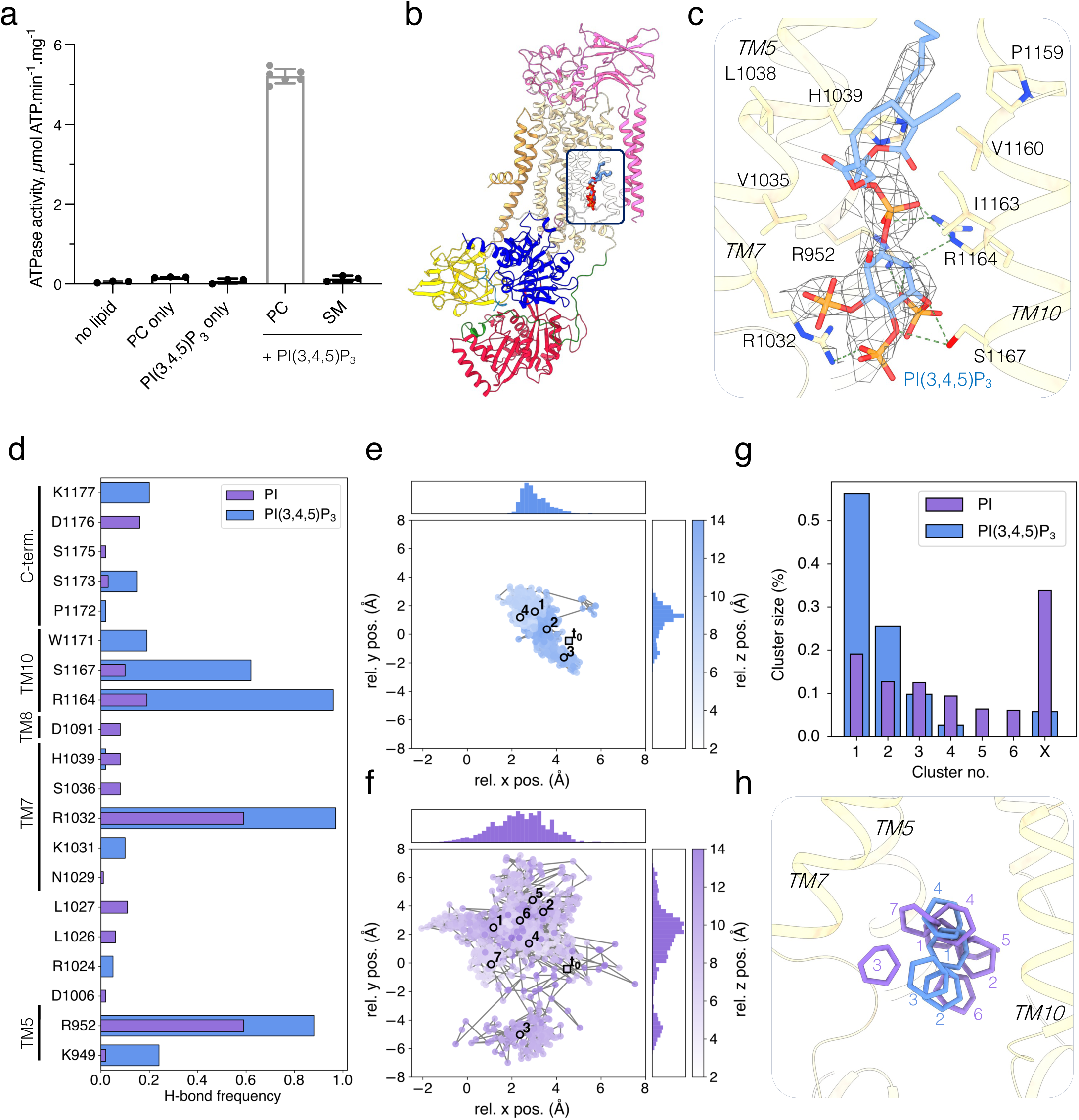
ATP8B1 PI(3,4,5)P_3_ binding site. **a)** ATPase activity of the C-terminally truncated ATP8B1-CDC50A complex in presence of PI(3,4,5)P_3_ and transport substrate (PC) at 37 °C. When annotated, the activity was measured in the presence of 1 mg·mL^-1^ DDM (2 mM), 0.17 mg·mL^-1^ PC or SM and a saturating concentration of 0.025 mg·mL^-1^ PI(3,4,5)P_3._ Data are mean ± s.d. of 3 replicated experiments (6 for PC + PI(3,4,5)P_3_) from the same purification. **b)** Overall structure of ATP8B1-CDC50A complex in the PI(3,4,5)P_3_ bound E2P_autoinhibited_ complex. Color code as Figure 1 with the N-terminal tail of ATP8B1 colored in blue. ATP8B1 rotated by 180° compared to overviews in Figure 1 and 2. **c)** Close-up view of the PI(3,4,5)P_3_ binding site and its associated EM density **d)** Hydrogen bonding (H-bond) frequencies between protein and the PI(3,4,5)P_3_ (blue) and PI (purple) headgroup from the last 200 ns of five MD trajectories of each system. **e-f)** Relative x, y and z positions of the center of mass (COM) of the inositide ring to the COM of selected Cɑ atoms of cavity-lining protein residues used as reference (see panel h) showing different mobility of the PI(3,4,5)P_3_ (blue; **e**) and PI (purple; **f**) headgroups. The highlighted points and numbers correspond to the cluster centroids in panel **h** and **t_0_** indicates the initial relative position of PI(3,4,5)P_3_ or PI according to the E2P_autoinhibited_ closed model. **g)** Clustering of inositol ring coordinates from the last 200 ns of each trajectory for PI(3,4,5)P_3_ (blue) and PI (purple). **h)** Structural visualization of the cluster centroids from clustering PI(3,4,5)P_3_ (blue) and PI (purple) inositol poses in the binding cavity (wheat). Only centroids from clusters containing ≥5% of the clustered structures are shown.

### The transport site of ATP8B1 can accommodate a wide range of glycerophospholipids

The transport substrate lipid POPC observed in the E2P_active_ structure adopts a similar conformation as observed for other substrate-bound P4-ATPases^26,31^. The PC headgroup is positioned at the end of a groove formed by TM1, 2,3 and 4 (Figure 3 & 5a). The acyl chains of the lipid are not interacting with the protein. Strikingly, the cavity surrounding the choline moiety formed by residues from TM1 and TM2 is wider than in other P4-ATPases characterized so far (Figure 5a & S9). We therefore hypothesized that with such a cavity, ATP8B1 should be able to bind a broad range of phospholipid headgroups. To test this hypothesis, we evaluated the ability of the main phospholipids found in eukaryotic membranes to stimulate the ATPase activity of ATP8B1 (Figure 5b). C-terminally truncated ATP8B1 was incubated with increasing lipid/DDM ratios, and a saturating concentration of PI(3,4,5)P_3_. All tested glycerophospholipids stimulated ATP8B1 ATPase activity, whereas SM did not (Figure 5b). Lipids with smaller head groups (PE, PG, PC or PA) seemed to be more activating than the negatively charged PS or the bulky PI. To better understand how ATP8B1 can accommodate such different lipid substrates, we determined the structure of the state associated with lipid occlusion and dephosphorylation. For this purpose, we used vanadate as an inhibitor that mimics the transition state of dephosphorylation of the catalytic aspartate, most likely by adopting a VO_3_^-^ planar conformation mimicking the leaving phosphate^32^. In this E2┅Pi like conformation, TM1 and 2 now enclose the substrate lipid, blocking the entry groove to prevent the entry of a new lipid (Figure 3). The A-domain has rotated and the catalytic E234 residue, from the conserved DGET motif of the A-domain, now coordinates a water molecule for in-line attack of the aspartyl phosphate moiety at the P-domain (Figure 1). We collected cryo-EM data sets in the presence of three lipid substrates spanning different chemical properties: PC as a standard lipid (Figure 5c), a negatively charged PS (Figure 5d), and a bulky PI (Figure 5e). The analysis of the lipid occlusion sites in each case revealed the presence of highly ordered water molecule networks interacting with the lipid headgroup and the protein (Figure 5f). The presence of this water network allows the interactions needed to close the groove formed in between TM 1, 2, 3 and 4 and the lipid headgroup. Interestingly, the structures showed that all three lipids are occluded in a similar manner, and side chains of the residues forming the occlusion site adopted virtually identical structures despite the presence of the charged carboxylic acid moiety of PS or the bulkier inositol moiety of PI (Figure 5g). However, the water molecule network differed significantly among the different lipid occlusion sites (Figure 5h). Our data suggest that lipid recognition, in the deepest part of the lipid groove triggers the organization of a water molecule network which seems to be part of the occlusion mechanism. This would be the triggering event of the allosteric transmission through TM2 that enables the A-domain rotation required for the dephosphorylation event. The differences observed between the organisation of water networks formed in presence of different lipids might explain the differences observed when comparing the ATPase activities.

**Figure 5.**
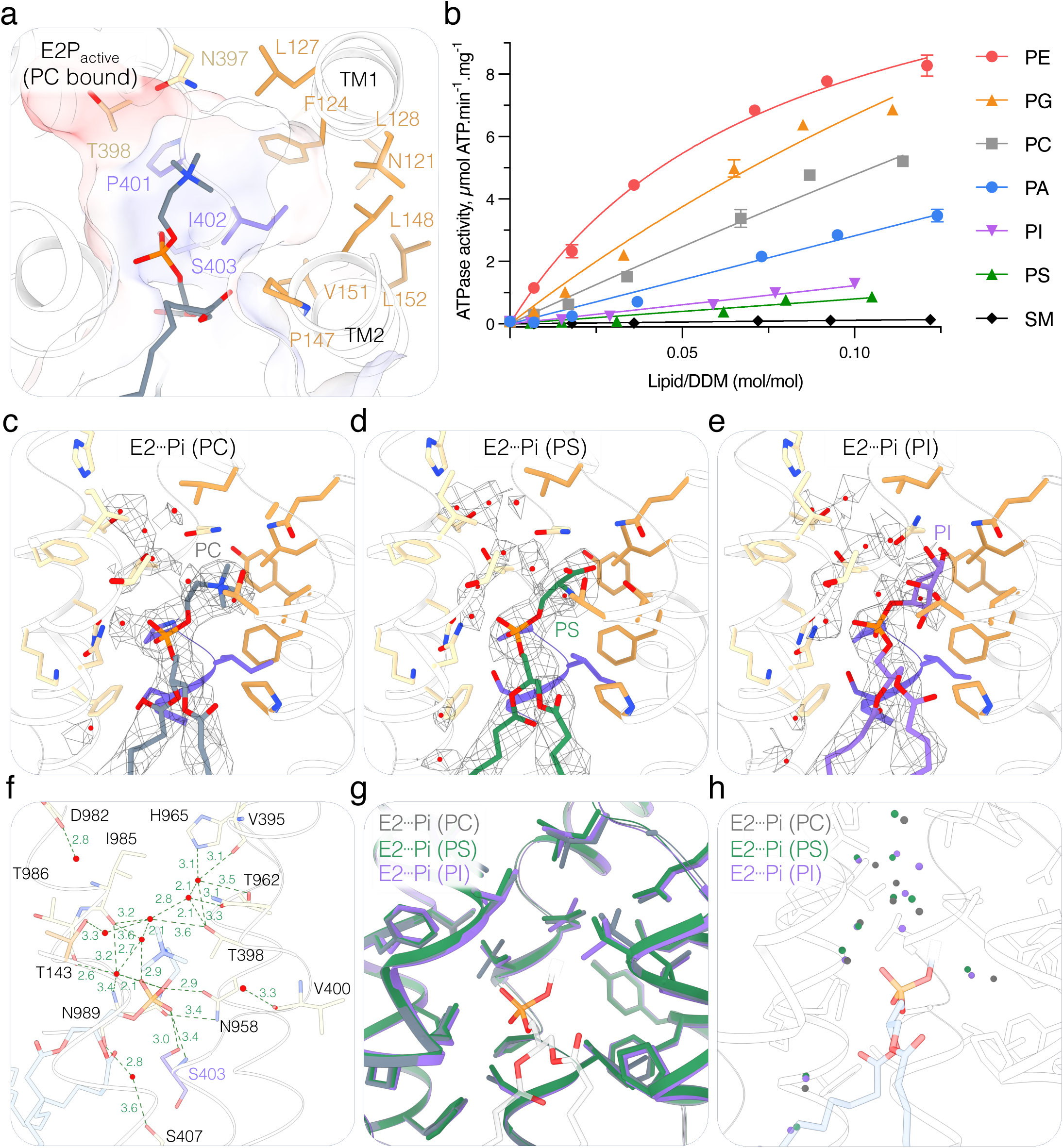
ATP8B1 lipid substrate specificity, binding and occlusion. **a)** Close-up view of the lipid binding site of ATP8B1 in the E2P_active_ state occupied by a PC molecule. The cavity formed by the protein around the lipid headgroup is shown as electrostatic surface. **b)** ATPase activity of the C-terminally truncated ATP8B1-CDC50A complex in presence of various phospholipids. The activity is measured at 37 °C with different lipid concentration but in presence of an invariant concentration of 1 mg·mL^-1^ DDM (2 mM) and 0.025 mg·mL^-1^ PI(3,4,5)P_3_. Data are mean ± s.d. of 3 replicates experiments from the same purification. **c-e)** Close up view of the lipid occlusion site of ATP8B1 in the E2┅Pi states occupied by PC **(c)**, PS **(d)** or PI **(e)**. The lipid associated EM density is shown as mesh. **f)** Detailed view of the water molecule network observed in lipid occlusion site of the E2┅Pi (PC) structure of ATP8B1-CDC50A complex. **g)** Cartoon representation of the structural alignment of the lipid occlusion site observed in the different E2┅Pi structure obtained with PC (grey), PS (green) or PI (purple). The PS molecule is shown without its head group to underline the lipid position in the three E2┅Pi structures. **h)** Comparison of the position of the water molecules observed in the lipid occlusion site.

### Sn-2 ester moiety is critical for substrate recognition

Given that all major glycerophospholipids tested were able to stimulate ATPase activity, unlike sphingomyelin, and can be recognized and occluded by the ATP8B1-CDC50A, we then decided to identify which part of the lipid is essential for substrate recognition. In the outward-open E2P_active_ state, the phosphoglycerol backbone of PC was found to interact directly with the PISL motif of ATP8B1. More specifically, the phosphate and the carbonyl function of the sn-2 ester bond interact via hydrogen bonding with the different rotamers of S403 (Figure 6a). Of note, in this conformation, the water molecule observed in the E1-ATP, E1P-ADP, E1P and the two E2P autoinhibited states is still at the same position, also interacting with the Ser403 side chain. For the E2┅Pi occluded state, however, the lipid is bound deeper in the pocket and the water molecule now displaced (Figure 6b). The different rotamers of the Ser403 now interact with the phosphate or with the sn-2 ester carbonyl more tightly (Figure 6b). To evaluate the importance of the sn-2 ester carbonyl interaction, we tested the ability of lipid analogs to stimulate the ATPase activity of the ATP8B1-CDC50A complex (Figure 6c). All lipids tested share the same choline headgroup and only differ in their chemical linkage between the phosphate group and the acyl chains. Plasmanylcholine, which contains an ether bond in sn-1 and an ester bond in sn-2, appeared to stimulate the ATPase activity almost as efficiently as PC (ester-PC). In contrast, ether-PC, which lacks the fatty acid in sn-2 position, failed to efficiently activate ATP8B1, achieving only 10% of the activity observed in presence of PC at equivalent lipid/DDM ratio. As previously described, lyso-PC, which lacks the fatty acid in sn-2 position can stimulate the ATPase activity but to a much lower extend than PC. Together, our data reveal that the ester bond in the sn-2 position is a critical determinant of lipid recognition by ATP8B1.

**Figure 6.**
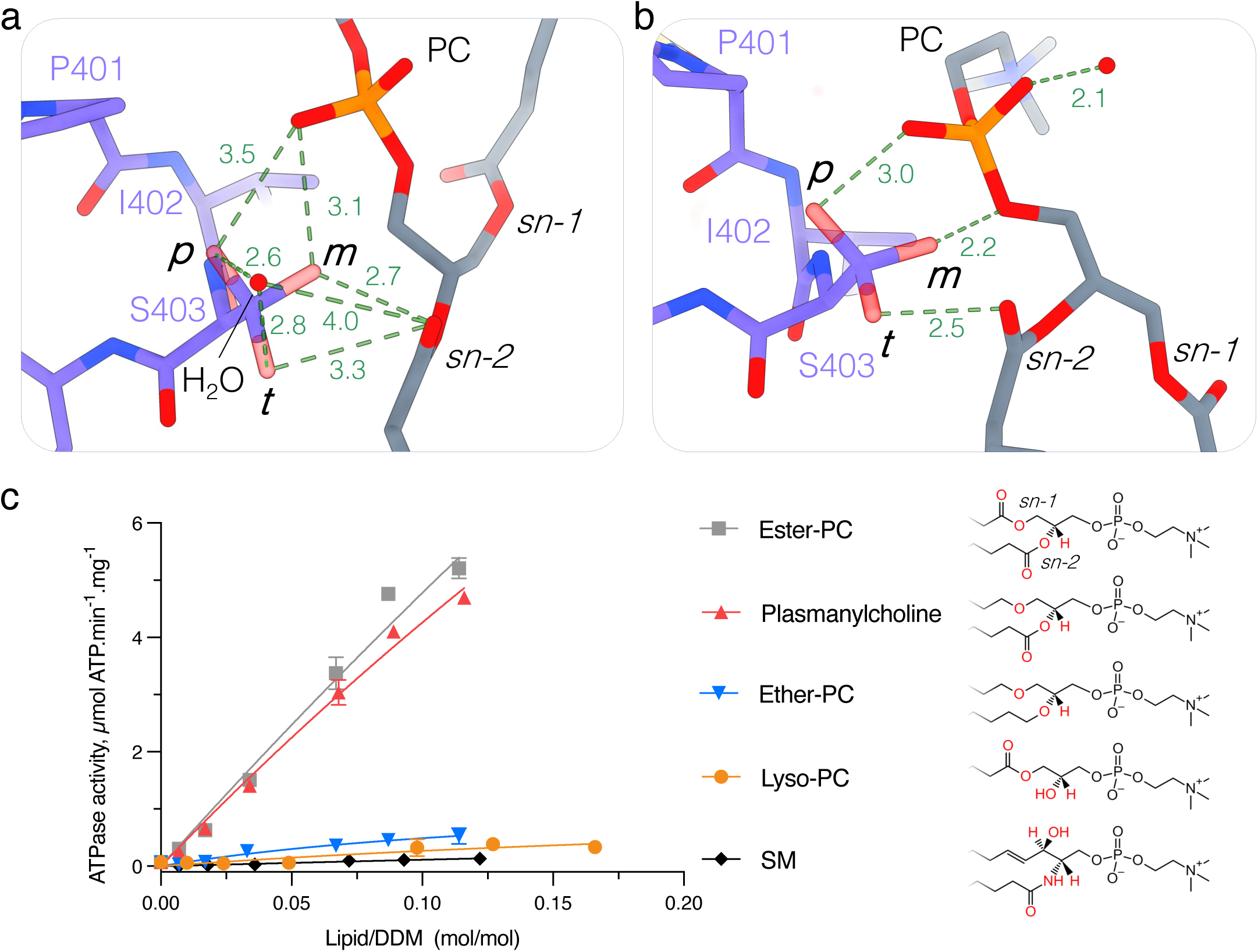
The sn-2 ester bond of the transport lipid is essential for substrate recognition. **a-b)** Close-up view of lipid and water interactions with the S403 of the PISL group of TM4 in the E2P_active_ (a) and E2┅Pi (PC) (b) state. The different rotamers detected by Coot of S403 are designated as p, m and t with a likelihood of occurrence of 48%, 29% and 22% respectively. **c)** ATPase activity of the C-terminally truncated ATP8B1-CDC50A complex in presence of different phospholipids with choline headgroup. The activity is measured at 37 °C with different lipid concentrations but in presence of an invariant concentration of 1 mg·mL^-1^ DDM (2 mM) and 0.025 mg·mL^-1^ PI(3,4,5)P_3_. Data are mean ± s.d. of 3 replicate experiments from the same purification.

## Discussion

### A water molecule at the transport site in P4- and P5-ATPases

Biochemical and structural studies of P4-ATPases have shown substrate-independent phosphorylation from ATP unlike for cation pumps where phosphorylation is tightly coupled to cation binding and occlusion. Thorough characterization of bovine ATP8A2-CDC50A and yeast Drs2p-Cdc50p also showed that phosphorylation was not dependent on a range of cations and quite insensitive to pH^11,33^. Lipid transport of ATP8A2 on solid-support membranes revealed a net outward charge transfer upon PS transport, also supporting substrate-independent phosphorylation^34,35^. Our studies agree with this model, although with the addition of a neutral water molecule in the transport site of P4-ATPases, which interacts with polar side chains in the absence of a substrate lipid headgroup (Figures 2c, S4). Interestingly, the water molecule stays bound in the outward-open E2P_active_ form of ATP8B1, where an incoming substrate lipid make first interactions with the lipid binding site, but the water molecule is chased upon occlusion of the substrate lipid, where the lipid headgroup directly interacts with the S403 side chain of the PISL motif, i.e. replacing the water molecule. This could be an important balance of water interaction versus lipid occlusion that fine-tunes the selection of proper lipid substrates. The role, pathways and dynamics of the water structure and how it is involved in the P4-ATPase cycle invite further studies by e.g. MD simulations that go beyond the scope of this paper. Interestingly, the structural and functional data obtained on P5A- and P5B-ATPases such as human ATP13A2 also show that substrate transport (of transmembrane alpha helices and polyamines, respectively) is coupled to dephosphorylation, whereas phosphorylation is independent of other substrates, i.e. similar to P4-ATPases. Congruent with this, the polyamine transport site of human ATP13A2 also binds water molecules at the transport site in their E1 to E1P half-cycle^27^ (Figure S5).

### Autoinhibition allows lipid binding, but not occlusion

Regulation through N- and C-terminal tails is a feature shared among various P-type ATPases, but structurally it was first described for the Drs2p-Cdc50p flippase complex, which revealed how the C-terminal tail occupies the nucleotide binding pocket and locks the cytoplasmic domains mouvements^28^. ATP8A1 and ATP8B1 structures showed a similar mechanism for ATP8-type lipid flippases in general^7,23,24^. C-terminal autoregulation locks the A-domain and restricts TM1-2 movements required for the opening of the outward-oriented lipid entry pathway (Figure S7). We show here that despite the presence of the N- and C-terminal tails at the interface of the cytoplasmic domains, the A-domain maintains some level of movement and TM1-2 can rotate to open the lipid entry pathway. However, lipid occlusion and dephosphorylation cannot occur. Rather, our data indicate that the E2P_autoinhibited_ state of ATP8B1 explores both open and closed conformations, and the open state can proceed to lipid binding. In our experimental conditions (4 °C in LMNG/CHS/PI(3,4,5)P_3_ mixed micelles), the closed conformation seems more to be favored, but the equilibrium could be poised toward the open conformation in a native membrane. Further structural investigations of ATP8B1 in a more native environment will be needed to evaluate sampling between lipid-free and lipid binding conformations in the autoinhibited form and how lipid might trigger further activation.

### ATP8B1 activation by phosphoinositides

ATP8B1 is, to our knowledge, the first human transporter found to be directly activated by PI(3,4,5)P_3_. From the cryo-EM data we identified a PI(3,4,5)P_3_ binding site in a positively charged cavity lined by TM 8, 9 and 10. Additionally, the binding site geometry remains unchanged through all conformations of the catalytic cycle, suggesting that the regulatory lipid binds in a conformation-independent manner (Figure S10a). Our MD data shows a clear interaction and stabilization of PI(3,4,5)P_3_ compared to PI in the identified pocket. However, questions remain open on how phosphoinositides activate ATP8B1 and potentially interfere with autoinhibition, and which specific steps of the catalytic cycle are stimulated. For Drs2p-Cdc50, PI(4)P destabilizes binding of the autoinhibitory C-terminal tail and stimulates dephosphorylation of E2P coupled to lipid occlusion and transport^28,36^. We would expect similar stimulation of E2P dephosphorylation for ATP8B1. In support of this, the E1 to E2P transition, corresponding to the autophosphorylation part of the cycle, of purified ATP8B1-CDC50A is independent of phosphoinositides^23^. Our studies do not provide immediate clues on the direct link between phosphoinositide binding and activation of ATP8B1, but it is likely linked to substrate recognition and/or occulsion. In our study we could observe that PI(3,4,5)P_3_ interacts tighly with the R952, which is located two residues after K957 on TM5, the later being part of the lipid transport site, and in interaction with previously described water molecule (Figure S10b). Thus, future work should focus on the identification of the PI(3,4,5)P_3_ activation mechanism.

### ATP8B1 has a broad lipid specificity

There is thus far no consensus about the transport substrate of ATP8B1^17–24^. Our structures for the E2P_active_ state with PC, PS, and PI show that different lipid head groups are accommodated at the lipid binding site - and more importantly, they can be occluded. This strongly suggests that they are also transport substrates. In comparison, the lipid headgroup binding cavity of other P4-ATPases is more restrictive compared to ATP8B1 (Figure S9). A screen for glycerophospholipids showed that only PS and PE stimulated ATPase activity and were shown to be transported by ATP11C^21,31,37^. A sequence alignment of ATP8B1 to PS/PE-specific flippases shows a high degree of conservation at the lipid binding cavity (Figure S11). Only few residues differ significantly, and surprisingly, they are not located around the lipid headgroup. A comparison of ATP11C and ATP8B1 in lipid-bound E2P states (i.e. where lipid recognition takes place prior to occlusion) shows that TM1 and TM2 are oriented differently. In ATP11C, TM1-2 are closer to TM3-4, thus diminishing the size of the groove for the lipid binding (Figure S12). Interestingly, the W390 residue in TM4 of ATP8B1 differs from the residues of the PS-specific flippases, where it is typically a leucine. The bulky W390 side chain keeps TM1-2 at a further distance from TM3-4 and provides a wider and more permissive groove for lipid head group recognition and occlusion. This points to the phosphoglycerol backbone as the critical determinant for substrate specificity of ATP8B1. Previous studies on ATP8A1 showed that ether-PS is not transported across erythrocyte membranes^38^. Here, we provide evidence that the carbonyl group of the sn-2 ester bond plays a critical role in substrate induced ATPase activity in ATP8B1, most likely due to the interactions it forms with the phosphate moiety, S401 of the conserved PISL motif and the water network upon substrate recognition and occlusion (Figure 6a-b). Sphingosine based-lipids, like SM, are not substrates of ATP8B1. In sphingolipids, the sn-2 ester bond is replaced by an amide bond, which engages in intra-molecular hydrogen bonds with the phosphate group^39^. In our three different lipid-occluded E2┅Pi structures (mimicking dephosphorylating E2P), the lipid phosphate interacts simultaneously with S401 and a highly ordered water molecule (Figures 5c-e & 6b). Hence, the internal H-bond of SM would disfavor the interactions observed for glycerophospholipids. Thus, our study broadens the lipid specificity of ATP8B1 to *sn1*-mono-ether phospholipids which is in line with a recently identified link between ATP8B2 and plasmalogens homeostasis^40^.

## Conclusions

In summary, we present a series of nine cryo-EM structures of the ATP8B1-CDC50A complex with accompanying functional studies and MD simulations, deciphering most of its catalytic cycle. The structures highlight a so far unrecognized water molecule bound at the lipid transport site during the first part of the catalytic cycle. Additionally, we observe two different autoinhibited states, a closed and an outward-open conformation which are likely in a dynamic equilibrium. Finally, our work reveals important determinants of lipid specificity in P4-ATPases and explains earlier discrepancies about the ATP8B1 substrate. The mechanism of activation based on phosphoinositide binding and interactions of the N- and C-terminal tails seem to entails subtle, but important effects and require further investigation.

## Methods

### Co-expression of the hATP8B1-hCDC50A complex

hATP8B1 (Uniprot: O43520; A1152T natural variant) and hCDC50A (Uniprot: Q9NV96) were co-expressed in a protease deficient *Saccharomyces cerevisiae* W303.1b/*Δpep4* (*MATα, leu2-3, his3-11, ura3-1, ade2-1, Δpep4, can^r^, cir^+^*) yeast strain as previously described^23^. To generate a C-terminal cleavable construct, an overlap extension PCR strategy was used to insert 8 aminoacids forming a HRV 3C protease site (LEVLFQGP) in between R1185 and K1187 of ATP8B1 sequence.

### Purification of the ATP8B1-CDC50A complex

Yeast cells were resuspended in buffer A (50 mM Tris-HCl pH 7.5, 1 mM EDTA, 0.6 M sorbitol) supplemented with protease inhibitors (2 mM PMSF, 2 µg.mL^-1^ leupeptine, 2 µg.mL^-1^ pepstatin, 2 µg.mL^-1^ chymostatin). The cells were subsequently broken with 0.5 mm glass beads using bead beaters. The crude extract was then spun down at 3,000 *g* for 30 min at 6 °C, to remove cell debris and nuclei. The membrane fraction was pelleted at 100,000 *g* for 90 min at 4 °C. The resulting yeast membrane pellets were finally resuspended at about 50 mg.mL^-1^ of total protein in buffer A supplemented with protease inhibitors (1 mM PMSF, 1 µg.mL^-1^ leupeptine, 1 µg.mL^-1^ pepstatin, 1 µg.mL^-1^ chymostatin). Membranes were diluted to 5.5 mg.mL^-1^ of total protein in ice-cold buffer B (50 mM MOPS-Tris at pH 7, 100 mM KCl, 1 mM DTT, 20% (w/v) glycerol and 5 mM MgCl_2,_), supplemented with protease inhibitors (1 mM PMSF, 1 µg.mL^-1^ leupeptine, 1 µg.mL^-1^ pepstatin, 1 µg.mL^-1^ chymostatin). The suspension was then supplemented with 15 mg.mL^-1^ DDM and 5 mg.mL^-1^ CHS and incubated 1 h at 4 °C. Washed membranes were pelleted by centrifugation at 100,000 *g* for 1 h at 6 °C. Insoluble material was pelleted by centrifugation at 125,000 *g* for 1 h at 4 °C. The supernatant, containing solubilized proteins, was applied onto a streptavidin-sepharose resin and incubated for 2 h at 6 °C. The resin was washed twice with six resin volumes of ice-cold buffer B supplemented with 0.2 mg.mL^-1^ LMNG and 0.02 mg.mL^-1^ CHS and protease inhibitors (1 mM PMSF, 1 µg.mL^-1^ leupeptine, 1 µg.mL^-1^ pepstatin, 1 µg.mL^-1^ chymostatin). The resin was then washed thrice with six resin volumes of ice-cold buffer B supplemented with 0.1 mg.mL^-1^ LMNG and 0.01 mg.mL^-1^ CHS. Elution was performed by the addition of 60 µg of purified TEV protease per mL of resin and overnight incubation at 6 °C. The eluted fraction was concentrated using a Vivaspin unit (100 kDa MWCO) prior to injection on a size-exclusion Superose 6 10/300GL increase column equilibrated with buffer C (50 mM MOPS-Tris pH 7, 100 mM KCl, 1 mM DTT, 5 mM MgCl_2_) supplemented with 0.03 mg.mL^-1^ LMNG and 0.003 mg.mL^-1^ CHS. For 3C protease site-containing version of ATP8B1, 100 µg of purified 3C protease (1:20 mol:mol ATP8B1:3C protease) were added to the concentrated sample and incubated 18h at 6 °C and 1h at 30 °C prior to size-exclusion chromatography.

### Cryo-EM sample preparation

Different conformations of the ATP8B1-CDC50A complex were stabilized in the following way: For the determination of the E1-ATP (AMPPCP), E1P-ADP (ADP+AlF_4_^-^), E1P (AlF_4_^-^), E2┅Pi [PC] (VO_4_ + POPC), E2┅Pi [PS] (VO_4_ + POPS) and E2┅Pi [PI] (VO_4_ + soy PI) states, the C-terminally truncated construct was incubated with 0.05 mg.mL^-1^ PI(4,5)P_2_ during 3C protease treatment. The sample was then injected on a size-exclusion Superose 6 10/300GL increase column equilibrated with buffer C supplemented with 0.03 mg.mL^-1^ LMNG and 0.003 mg.mL^-1^ CHS. The purified complex was then concentrated to 0.8-5 mg.mL^-1^ and incubated with PI(4,5)P_2_ 0.0015 mg.mL^-1^ and respectively with 1mM AMPPCP / 3 mM ADP + 2 mM AlCl_3_ + 10 mM NaF / 2 mM AlCl_3_ + 10 mM NaF / 1 mM Na_3_VO_4_ + 0.003 mg.mL^-1^ POPC / 1 mM Na_3_VO4 + 0.003 mg.mL^-1^ POPS / 1 mM Na_3_VO4 + 0.003 mg.mL^-1^ soy PI for 1 h on ice.For the E2P_active_ state: the C-terminally truncated construct was incubated with 0.01 mg.mL^-1^ POPC and 0.002 mg.mL^-1^ PI(3,4,5)P_3_ during elution from the streptavidin resin by TEV protease. The sample was then injected on a size-exclusion Superose 6 10/300GL increase column equilibrated with buffer C supplemented with 0.003 mg.mL^-1^ POPC and 0.0015 mg.mL^-1^ PI(3,4,5)P_3_. The purified complex was then concentrated to 5 mg.mL^-1^ and incubated with 1 mM BeSO_4_ and 4 mM KF for 1 h on ice. For the E2P_autoinhibited_ states: the WT full-length construct was incubated with 0.002 mg.mL^-1^ PI(3,4,5)P_3_ during elution form the streptavidin resin by TEV. After concentration the sample was incubated with 2 mM ATP. The sample was then injected on a size-exclusion Superose 6 10/300GL increase column equilibrated with buffer C supplemented with 0.0015 mg.mL^-1^ PI(3,4,5)P_3_. For this sample CHS was omitted from all steps of the purification as we initially hypothesized that CHS might compete with PI(3,4,5)P_3_. The purified complex was then concentrated to 3 mg.mL^-1^. In all cases, 3 µL of the final sample were added to freshly glow-discharged (45 seconds at 15 mA) C-flat Holey Carbon grids, CF-1.2/1.3-4C (Protochips), which were subsequently vitrified at 4 °C and 100% humidity on a Vitrobot IV (Thermo Fisher Scientific).

### Cryo-EM data collection

All data were collected on a Titan Krios G3i (EMBION Danish National cryo-EM Facility – Aarhus node) with X-FEG operated at 300 kV and equipped with a Gatan K3 camera and a Bioquantum energy filter using a slit width of 20 eV. Movies were collected using aberration-free image shift data collection (AFIS) in EPU (Thermo Fisher Scientific) as 1.5 second exposures in super-resolution mode at a calibrated physical pixel size of 0.647 Å/pixel (magnification of 130,000x) with a total electron dose of 60 e^-^/Å^2^.

### Cryo-EM data processing

Processing was performed in cryoSPARC^41^ (v3). Patch Motion Correction and Patch CTF were performed before low-quality micrographs (e.g. micrographs with crystalline ice, high motion) were discarded. Particles were initially picked using template picking on all movies. Particles were extracted in a 416-pixel box and Fourier cropped to a 104-pixel box (2.59 Å/pixel). *Ab initio* references were produced using a subset of all particles selected from 2D class. One protein-like reference and multiple junk references were used in multiple rounds of heterogeneous refinement. Selected particles were then re-extracted in an appropriate pixel box depending of the maximum resolution achieved with non-uniform refinement with a first extraction in a 208-pixel box (1.3 Å/pixel). The final particle stack was used to perform a final non-uniform refinement combined with per-particle CTF refinement^42^. The resulting map was used to generate a mask to run 3D variability analysis to investigate sample flexibility and heterogeneity^43^. For the two E2P autoinhibited states, conformational heterogeneity was detected, and the two final particles stack were subjected to a final non-uniform refinement combined with per-particle CTF refinement.

### Model building

The ATP8B1-CDC50A models were built using the previously published structure of the complex (PDB: 7PY4) as templates. The cryo-EM maps were sharpened using the Autosharpen tool in PHENIX^44^. The models were manually generated and relevant ligands added with COOT^45^ before real space refinement in PHENIX^46^. For real space refinement, hydrogen addition and ligand restraints were generated by the ReadySet tool in PHENIX. Model validation was performed using MolProbity^47^ in PHENIX, and relevant metrics are listed in Table S1.

### Molecular dynamic simulation and analysis

The starting structures of the ATP8B1/CDC50A complex for the MD simulations were based on the cryo-EM model of the E2P_autoinhibited_ “closed” state (PDB:XXXX). We modelled missing loops (28-62, 72-105, 714-786 and 814-832) using MODELLER^48^ (v10.1). The template for loop 2 (72-105) was based on the corresponding AlphaFold2 model^49^ because of the presence of a local hairpin structure with reasonable confidence and confirmed by the density observed in the E2P active conformation. We modelled the phosphorylated D454 with two negative charges and parameterized as previously described^50^. We kept the Mg^2+^ ion and three disulfide bonds in CDC50A (C91-C104, C94-C102 and C157-C171) but removed glycans. E230 and E843 were protonated and K957 was deprotonated based on predictions using PROPKA3^51^ to match the experimental conditions at pH 7. The IP9 molecule in the cryo-EM structure was replaced by a PI or PI(3,4,5)P3, modelled as a SAPI (18:0/20:4) and SAPI34 (18:0/20:4), respectively. We set up the system using the CHARMM-GUI Membrane Builder web tool^52^ and parameterized the system with the Charmm36m force field^53^ in conjunction with the TIP3 water model^54^ using the CHARMM-GUI Input Generator^55^. The protein complex was embedded in a bilayer, and we chose the lipid composition of the cytosolic (inner/lower) leaflet and luminal (outer/upper) leaflet to mimic a mammalian plasma membrane^56,57^. The lower leaflet consists of 15% cholesterol (CHOL), 20% phosphatidylcholine (13% PLPC and 7% POPC), 33% phosphatidylethanolamine (YOPE), 23% phosphatidic acid (YOPA), 3% phosphatidylserine (POPS), and 6.0% phosphatidylinositol (3% SAPI, 1% SAPI14, 1% SAPI2D, and 1% SAPI34). The upper leaflet is composed of 39% cholesterol (CHOL), 46% phosphatidylcholine (31% PLPC and 15% POPC), and 15% sphingomyelin (SM). The membrane spans an area of 14 x 14 nm, totaling to about 300 lipids in each leaflet. We solvated the systems with about 92,000 water molecules, neutralized the systems and matched experimental ion concentrations of 100 mM KCl and 5 mM MgCl_2_ with corresponding ions. The total atom number was around 370,000 and the equilibrated box dimensions in the x, y and z dimension were 13.5 nm, 13.5 nm and 19.9 nm, respectively. We minimized the systems with 5,000 steps of steepest descent and positional restraints on protein backbone and side chain atoms and lipid atoms. We performed two steps of equilibration in the NVT ensemble (250 ps each) using the Berendsen thermostat^58^ and a constant temperature of 310 K. This was followed by 4 more steps of equilibration in the NPT ensemble of 500 ps each by introducing semi-isotropic pressure coupling using the Berendsen barostat. Positional restraints were gradually released during equilibration steps. We performed the production run in the NPT ensemble at 310 K using the Noose-Hoover thermostat and a coupling constant of 1 ps. The pressure was kept constant at 1 bar with semi isotropic pressure coupling using the Parrinello-Rahman barostat^59^, a 5 ps time coupling constant and an isothermal compressibility of 4.5.10^-5^ bar ^-1^. We applied a 1.2 nm cutoff for the van der Waals interactions using a switching function from 1.0 nm and a cutoff of 1.2 nm for short-range electrostatic interactions while long-range electrostatics were treated by Particle-mesh Ewald summation^60^. The Verlet algorithm was used to generate a pair list with buffer and the time step for integration was 2 fs. For each system (PI-bound and PI(3,4,5)P_3_-bound), we performed a set of five 600 ns long production runs. Finally, we analyzed the last 200 ns of each simulation, pooled the five replicates and used a frame every 1 ns. All simulations were run using GROMACS^61^ (v2020/3). Analysis was performed using GROMACS v 2019/4 and python using the libraries MDAnalysis and GetContacts. To calculate relative positions of the inositol ring, we rotated and translated the trajectory by aligning Cɑ atoms of TM residues in proximity to the lipid binding site and used the center of mass of the corresponding atoms as reference. The binding poses were clustered using the gromos algorithm^30^ based on the C atoms in the inositol ring using a cutoff of 0.1 nm.

### ATPase activity

The rate of ATP hydrolysis was monitored continuously using an enzyme-coupled assay, coupling ATP hydrolysis to a drop in NADH absorption at 340 nm. ATPase activity was measured at 37 °C in buffer C supplemented with 1 mM ATP, 1 mM phosphoenolpyruvate, 0.04 mg.mL^-1^ pyruvate kinase, 0.1 mg.mL^-1^ lactate dehydrogenase, 370 µM NADH, 1 mg.mL^-1^ DDM (2 mM), 0.025 mg.mL^-1^ PI(3,4,5)P_3_ and variable amounts of other phospholipids. In these experiments, the purified C-terminally truncated ATP8B1-CDC50A complex was added at a final concentration 2.3 µg.mL^-1^. The NADH absorption decay was measured in a 96-well plate with a SpectraMax®i3 microplate reader in kinetic mode. Conversion from NADH oxidation rates expressed in mAU.s^-1^ to ATPase activities expressed in µmol.min^-1^.mg^-1^ was based on the extinction coefficient of NADH at 340 nm.

## Acknowledgments

We thank Thomas Boesen, Andreas Bøggild and Taner Drace for technical support during EM data collection at the EMBION Danish National cryo-EM facility of Aarhus University (5072-00025B, Danish Agency for Research and Higher Education), Jesper Lykkegaard Karlsen for scientific computing support. We are grateful to Anna Marie Nielsen, Tanja Klymchuk and Bente Andersen (Aarhus University) for technical assistance. We also wish to thank, Tomás Heger, Line Marie Christiansen, Ronja Driller and David Stokes for discussion and advice.

This work was supported by a Marie Skłodowska-Curie Actions Individual Fellowship — LivFlip (101024542) to T.D., by Engineering and Physical Sciences Research Council grants (EP/X035603 and EP/V030779) to S. K., by an ANR grant (ANR-14-CE09-0022) to G.L., by the French Infrastructure for Integrated Structural Biology (FRISBI; ANR-10-INSB-05) to G.L., by the Centre National de la Recherche Scientifique (CNRS) to G.L., by the Lundbeck Foundation to the BRAINSTRUC structural biology initiative (155-2015-2666) to K.L.-L. and P.N., and by a Lundbeckfonden Professorship grant (R310-2018-3713) to P.N.

## Author contributions

T.D., and P.N. conceived the project. T.D., F.K., K.L-L. and P.N. designed research. T.D. and F.K. performed experiments. T.D., F.K., C.S., R.K.F., S.K. J.A.L., K.L-L. and P.N. analyzed data. T.D. drafted the manuscript. All authors contributed to the manuscript.

## Notes

### Competing Interest Statement

The authors have declared no competing interest.

### Summary of Updates

- In the abstract, the PI(3,4,5)P3 binding site was described as an electronegative pocket while being electropositive. - Some typos were corrected.

## Bibliography

1. Clarke, R. J., Hossain, K. R. & Cao, K. Physiological roles of transverse lipid asymmetry of animal membranes. Biochim. Biophys. Acta BBA - Biomembr. 1862, 183382 (2020).

2. Doktorova, M., Symons, J. L. & Levental, I. Structural and functional consequences of reversible lipid asymmetry in living membranes. Nat. Chem. Biol. 16, 1321–1330 (2020).

3. Andersen, J. P. et al. P4-ATPases as Phospholipid Flippases—Structure, Function, and Enigmas. Front. Physiol. 7, (2016).

4. Scortecci, J. F. et al. Cryo-EM structures of the ABCA4 importer reveal mechanisms underlying substrate binding and Stargardt disease. Nat. Commun. 12, (2021).

5. Palmgren, M. G. & Nissen, P. P-type ATPases. Annu. Rev. Biophys. 40, 243–266 (2011).

6. Vestergaard, A. L. et al. Critical roles of isoleucine-364 and adjacent residues in a hydrophobic gate control of phospholipid transport by the mammalian P4-ATPase ATP8A2. Proc. Natl. Acad. Sci. 111, E1334–E1343 (2014).

7. Hiraizumi, M., Yamashita, K., Nishizawa, T. & Nureki, O. Cryo-EM structures capture the transport cycle of the P4-ATPase flippase. Science 365, 1149–1155 (2019).

8. Albers, R. W. Biochemical Aspects of Active Transport. Annu. Rev. Biochem. 36, 727– 756 (1967).

9. Post, R. L., Hegyvary, C. & Kume, S. Activation by adenosine triphosphate in the phosphorylation kinetics of sodium and potassium ion transport adenosine triphosphatase. J. Biol. Chem. 247, 6530–6540 (1972).

10. Olesen, C., Sørensen, T. L.-M., Nielsen, R. C., Møller, J. V. & Nissen, P. Dephosphorylation of the Calcium Pump Coupled to Counterion Occlusion. Science 306, 2251–2255 (2004).

11. Coleman, J. A., Vestergaard, A. L., Molday, R. S., Vilsen, B. & Andersen, J. P. Critical role of a transmembrane lysine in aminophospholipid transport by mammalian photoreceptor P4-ATPase ATP8A2. Proc. Natl. Acad. Sci. U. S. A. 109, 1449–1454 (2012).

12. Timcenko, M. et al. Structural Basis of Substrate-Independent Phosphorylation in a P4-ATPase Lipid Flippase. J. Mol. Biol. 433, 167062 (2021).

13. Bull, L. N. et al. A gene encoding a P-type ATPase mutated in two forms of hereditary cholestasis. Nat. Genet. 18, 219–224 (1998).

14. Groen, A. et al. Complementary functions of the flippase ATP8B1 and the floppase ABCB4 in maintaining canalicular membrane integrity. Gastroenterology 141, 1927–1937.e1–4 (2011).

15. Dumitrescu, L. et al. Genetic variants and functional pathways associated with resilience to Alzheimer’s disease. Brain 143, 2561–2575 (2020).

16. Holstege, H. et al. Exome sequencing identifies rare damaging variants in ATP8B4 and ABCA1 as risk factors for Alzheimer’s disease. Nat. Genet. 54, 1786–1794 (2022).

17. Paulusma, C. C. et al. ATP8B1 requires an accessory protein for endoplasmic reticulum exit and plasma membrane lipid flippase activity. Hepatol. Baltim. Md 47, 268–278 (2008).

18. Muñoz-Martínez, F., Torres, C., Castanys, S. & Gamarro, F. CDC50A plays a key role in the uptake of the anticancer drug perifosine in human carcinoma cells. Biochem. Pharmacol. 80, 793–800 (2010).

19. Ray, N. B. et al. Dynamic regulation of cardiolipin by the lipid pump Atp8b1 determines the severity of lung injury in experimental pneumonia. Nat. Med. 16, 1120–1127 (2010).

20. Takatsu, H. et al. Phospholipid flippase activities and substrate specificities of human type IV P-type ATPases localized to the plasma membrane. J. Biol. Chem. 289, 33543–33556 (2014).

21. Segawa, K., Kurata, S. & Nagata, S. Human Type IV P-type ATPases That Work as Plasma Membrane Phospholipid Flippases and Their Regulation by Caspase and Calcium. J. Biol. Chem. 291, 762–772 (2016).

22. Mizutani, A. et al. Assessment of Adenosine Triphosphatase Phospholipid Transporting 8B1 (ATP8B1) Function in Patients With Cholestasis With ATP8B1 Deficiency by Using Peripheral Blood Monocyte-Derived Macrophages. Hepatol. Commun. 5, 52–62 (2021).

23. Dieudonné, T. et al. Autoinhibition and regulation by phosphoinositides of ATP8B1, a human lipid flippase associated with intrahepatic cholestatic disorders. eLife 11, e75272 (2022).

24. Cheng, M.-T. et al. Structural insights into the activation of autoinhibited human lipid flippase ATP8B1 upon substrate binding. Proc. Natl. Acad. Sci. U. S. A. 119, e2118656119 (2022).

25. Jidenko, M., Lenoir, G., Fuentes, J. M., le Maire, M. & Jaxel, C. Expression in yeast and purification of a membrane protein, SERCA1a, using a biotinylated acceptor domain. Protein Expr. Purif. 48, 32–42 (2006).

26. Nakanishi, H. et al. Transport Cycle of Plasma Membrane Flippase ATP11C by Cryo-EM. Cell Rep. 32, 108208 (2020).

27. Sim, S. I., von Bülow, S., Hummer, G. & Park, E. Structural basis of polyamine transport by human ATP13A2 (PARK9). Mol. Cell 81, 4635–4649.e8 (2021).

28. Timcenko, M. et al. Structure and autoregulation of a P4-ATPase lipid flippase. Nature 571, 366–370 (2019).

29. Xu, J., He, Y., Wu, X. & Li, L. Conformational changes of a phosphatidylcholine flippase in lipid membranes. Cell Rep. 38, 110518 (2022).

30. Daura, X., et al. Peptide Folding: When Simulation Meets Experiment. Angew. Chem. Int. Ed. 38, 236–240 (1999).

31. Nakanishi, H. et al. Crystal structure of a human plasma membrane phospholipid flippase. J. Biol. Chem. 295, 10180–10194 (2020).

32. Clausen, J. D. et al. Crystal Structure of the Vanadate-Inhibited Ca(2+)-ATPase. Struct. Lond. Engl. 1993 24, 617–623 (2016).

33. Jacquot, A. et al. Phosphatidylserine stimulation of Drs2p·Cdc50p lipid translocase dephosphorylation is controlled by phosphatidylinositol-4-phosphate. J. Biol. Chem. 287, 13249–13261 (2012).

34. Tadini-Buoninsegni, F., Mikkelsen, S. A., Mogensen, L. S., Molday, R. S. & Andersen, J. P. Phosphatidylserine flipping by the P4-ATPase ATP8A2 is electrogenic. Proc. Natl. Acad. Sci. U. S. A. 116, 16332–16337 (2019).

35. Tadini-Buoninsegni, F. et al. Electrogenic reaction step and phospholipid translocation pathway of the mammalian P4-ATPase ATP8A2. FEBS Lett. 597, 495–503 (2023).

36. Azouaoui, H. et al. Coordinated Overexpression in Yeast of a P4-ATPase and Its Associated Cdc50 Subunit: The Case of the Drs2p/Cdc50p Lipid Flippase Complex. in P-Type ATPases: Methods and Protocols (ed. Bublitz, M.) 37–55 (Springer, 2016). doi:10.1007/978-1-4939-3179-8_6.

37. Wang, J. et al. Proteomic Analysis and Functional Characterization of P4-ATPase Phospholipid Flippases from Murine Tissues. Sci. Rep. 8, 10795 (2018).

38. Fellmann, P. et al. Transmembrane Movement of Diether Phospholipids in Human Erythrocytes and Human Fibroblasts. Biochemistry 39, 4994–5003 (2000).

39. Slotte, J. P. The importance of hydrogen bonding in sphingomyelin’s membrane interactions with co-lipids. Biochim. Biophys. Acta BBA - Biomembr. 1858, 304–310 (2016).

40. Honsho, M., Mawatari, S. & Fujiki, Y. ATP8B2-Mediated Asymmetric Distribution of Plasmalogens Regulates Plasmalogen Homeostasis and Plays a Role in Intracellular Signaling. Front. Mol. Biosci. 9, 915457 (2022).

41. Punjani, A., Rubinstein, J. L., Fleet, D. J. & Brubaker, M. A. cryoSPARC: algorithms for rapid unsupervised cryo-EM structure determination. Nat. Methods 14, 290–296 (2017).

42. Punjani, A., Zhang, H. & Fleet, D. J. Non-uniform refinement: adaptive regularization improves single-particle cryo-EM reconstruction. Nat. Methods 17, 1214–1221 (2020).

43. Punjani, A. & Fleet, D. J. 3D variability analysis: Resolving continuous flexibility and discrete heterogeneity from single particle cryo-EM. J. Struct. Biol. 213, 107702 (2021).

44. Terwilliger, T. C., Sobolev, O. V., Afonine, P. V. & Adams, P. D. Automated map sharpening by maximization of detail and connectivity. Acta Crystallogr. Sect. Struct. Biol. 74, 545–559 (2018).

45. Emsley, P., Lohkamp, B., Scott, W. G. & Cowtan, K. Features and development of Coot. Acta Crystallogr. D Biol. Crystallogr. 66, 486–501 (2010).

46. Afonine, P. V. et al. Real-space refinement in PHENIX for cryo-EM and crystallography. Acta Crystallogr. Sect. Struct. Biol. 74, 531–544 (2018).

47. Chen, V. B. et al. MolProbity: all-atom structure validation for macromolecular crystallography. Acta Crystallogr. D Biol. Crystallogr. 66, 12–21 (2010).

48. Webb, B. & Sali, A. Comparative Protein Structure Modeling Using MODELLER. Curr. Protoc. Bioinforma. 54, 5.6.1–5.6.37 (2016).

49. Jumper, J. et al. Highly accurate protein structure prediction with AlphaFold. Nature 596, 583–589 (2021).

50. Das, A., Rui, H., Nakamoto, R. & Roux, B. Conformational Transitions and Alternating-Access Mechanism in the Sarcoplasmic Reticulum Calcium Pump. J. Mol. Biol. 429, 647–666 (2017).

51. Olsson, M. H. M., Søndergaard, C. R., Rostkowski, M. & Jensen, J. H. PROPKA3: Consistent Treatment of Internal and Surface Residues in Empirical pKa Predictions. J. Chem. Theory Comput. 7, 525–537 (2011).

52. Wu, E. L. et al. CHARMM-GUI Membrane Builder toward realistic biological membrane simulations. J. Comput. Chem. 35, 1997–2004 (2014).

53. Huang, J. et al. CHARMM36m: an improved force field for folded and intrinsically disordered proteins. Nat. Methods 14, 71–73 (2017).

54. Jorgensen, W. L., Chandrasekhar, J., Madura, J. D., Impey, R. W. & Klein, M. L. Comparison of simple potential functions for simulating liquid water. J. Chem. Phys. 79, 926–935 (1983).

55. Lee, J. et al. CHARMM-GUI Input Generator for NAMD, GROMACS, AMBER, OpenMM, and CHARMM/OpenMM Simulations Using the CHARMM36 Additive Force Field. J. Chem. Theory Comput. 12, 405–413 (2016).

56. Lorent, J. H. et al. Plasma membranes are asymmetric in lipid unsaturation, packing and protein shape. Nat. Chem. Biol. 16, 644–652 (2020).

57. van Meer, G., Voelker, D. R. & Feigenson, G. W. Membrane lipids: where they are and how they behave. Nat. Rev. Mol. Cell Biol. 9, 112–124 (2008).

58. Berendsen, H. J. C., Postma, J. P. M., van Gunsteren, W. F., DiNola, A. & Haak, J. R. Molecular dynamics with coupling to an external bath. J. Chem. Phys. 81, 3684–3690 (1984).

59. Parrinello, M. & Rahman, A. Polymorphic transitions in single crystals: A new molecular dynamics method. J. Appl. Phys. 52, 7182–7190 (1981).

60. Essmann, U. et al. A smooth particle mesh Ewald method. J. Chem. Phys. 103, 8577– 8593 (1995).

61. Abraham, M. J. et al. GROMACS: High performance molecular simulations through multi-level parallelism from laptops to supercomputers. SoftwareX 1–2, 19–25 (2015).

