## Supplementary data for "Activation and substrate specificity of the human P4-ATPase ATP8B1"

a

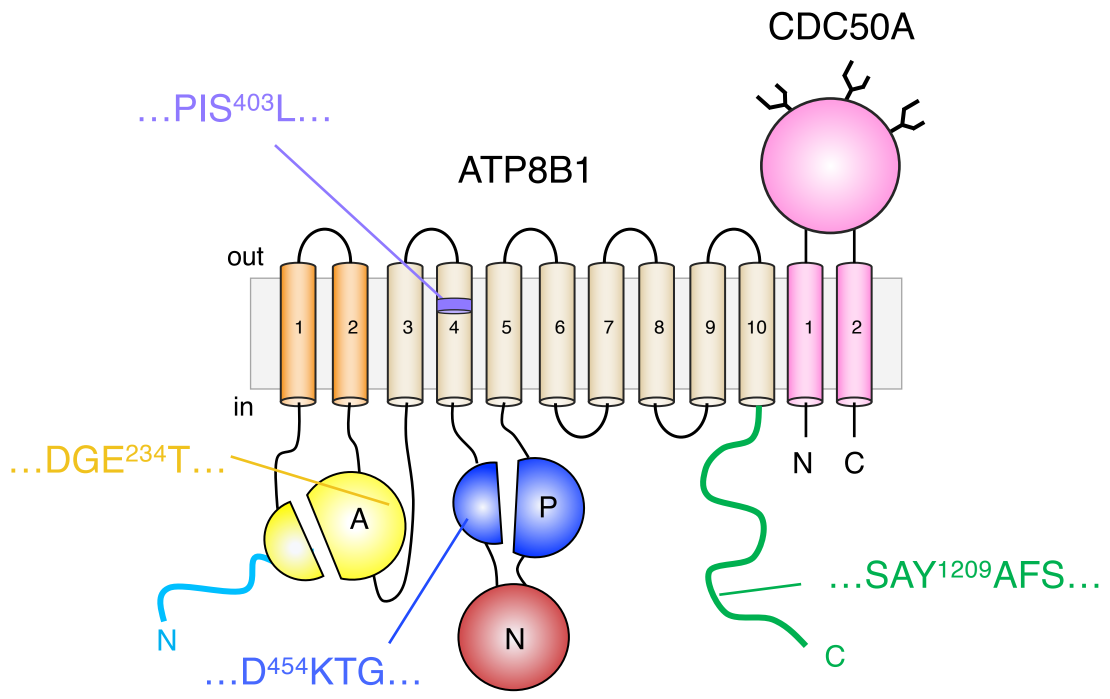


b

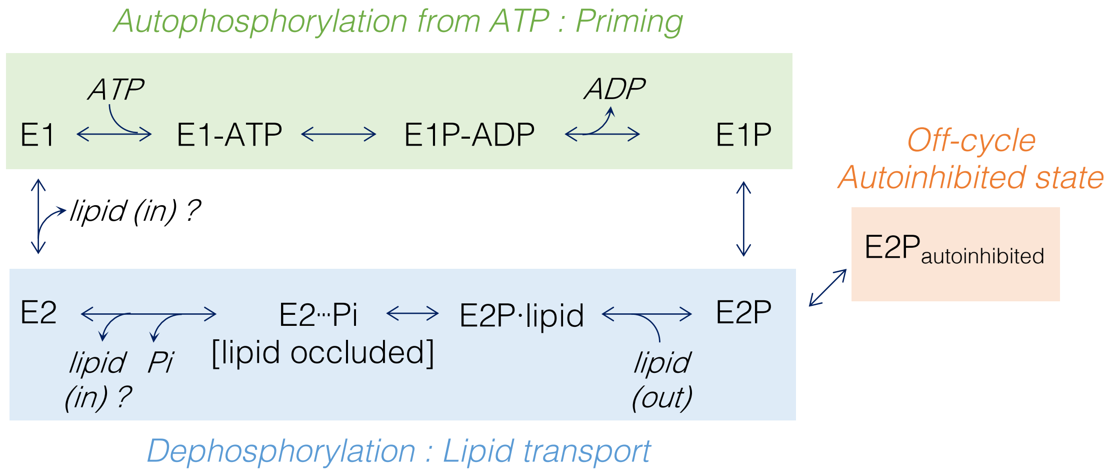


**Figure S1 – ATP8B1-CDC50A topology and Post-Albers P4-ATPase catalytic cycle**

**a)** The ATP8B1-CDC50A complex topology shows the Actuator domain (A), the Nucleotide binding domain (N), and the Phosphorylation domain (P) highlighted in yellow, red, and blue, respectively. The regulatory N- and C-terminal tails are shown in cyan and green, respectively. The P4-ATPase canonical motif of the phosphorylation site (DKTG, P domain), dephosphorylation loop (DGET, A domain), the transport site (PISL, TM4), and the autoinhibition signature motif (SAYAFS, C-terminal tail) are highlighted with ATP8B1 residue numbers. **b)** The Post-Albers P4-ATPase catalytic cycle includes the substrate-independent phosphorylation step from ATP (Priming) and the substrate-dependent dephosphorylation (transport). For autoinhibited P4-ATPases, an off-cycle E2P state prevents transport.

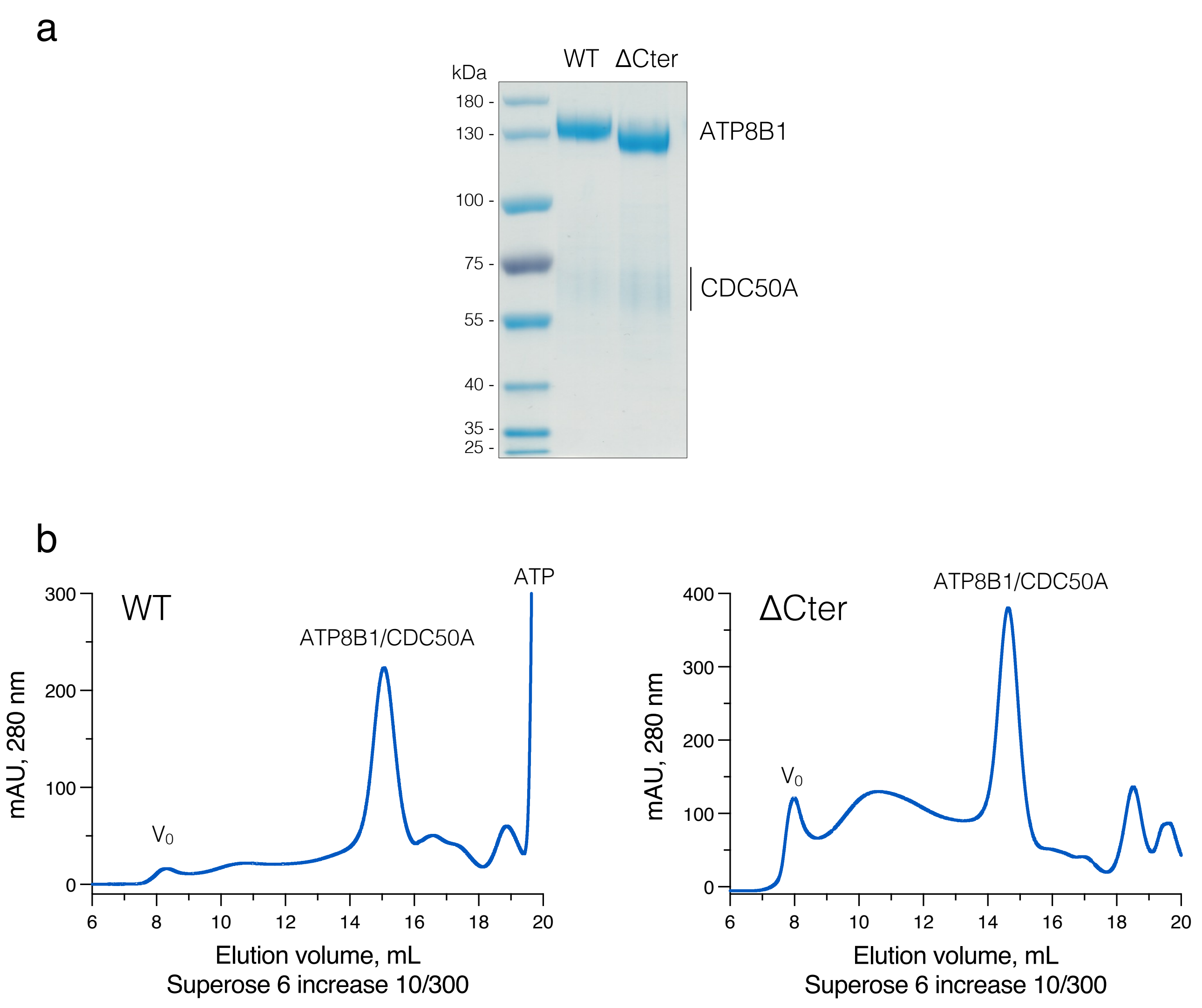


**Figure S2 – Biochemical characterization of the purified ATP8B1-CDC50A complex**

**a)** SDS-PAGE of the purified ATP8B1-CDC50A complexes (WT full length and C-terminally truncated) used for Cryo-EM and ATPase activity measurement in this study. **b)** Size exclusion chromatography (Superose 6 increase 10/300) elution profiles from the last purification step of the WT full length and C-terminally truncated ATP8B1-CDC50A complexes.

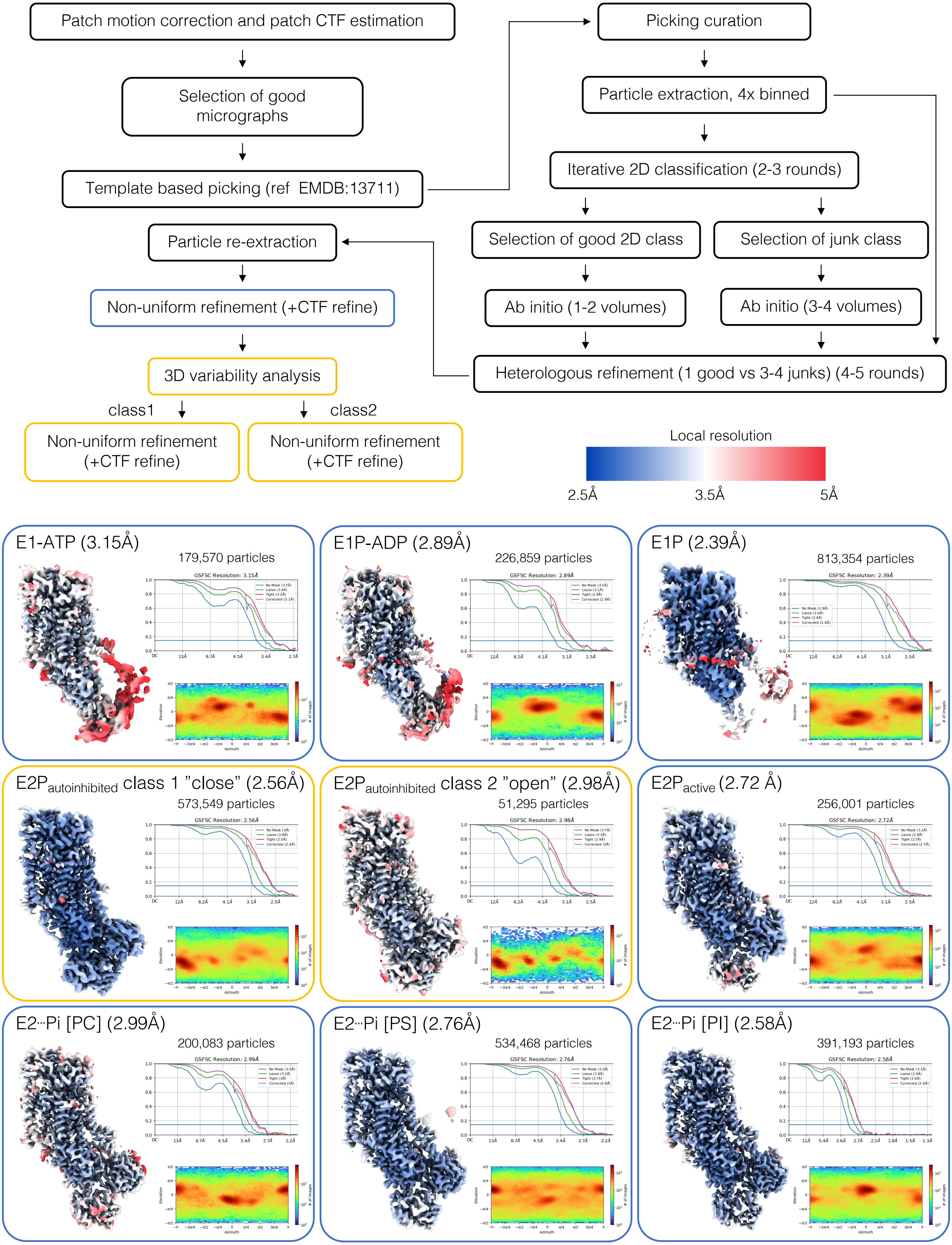


**Figure S3 – Cryo-EM data processing pipeline**

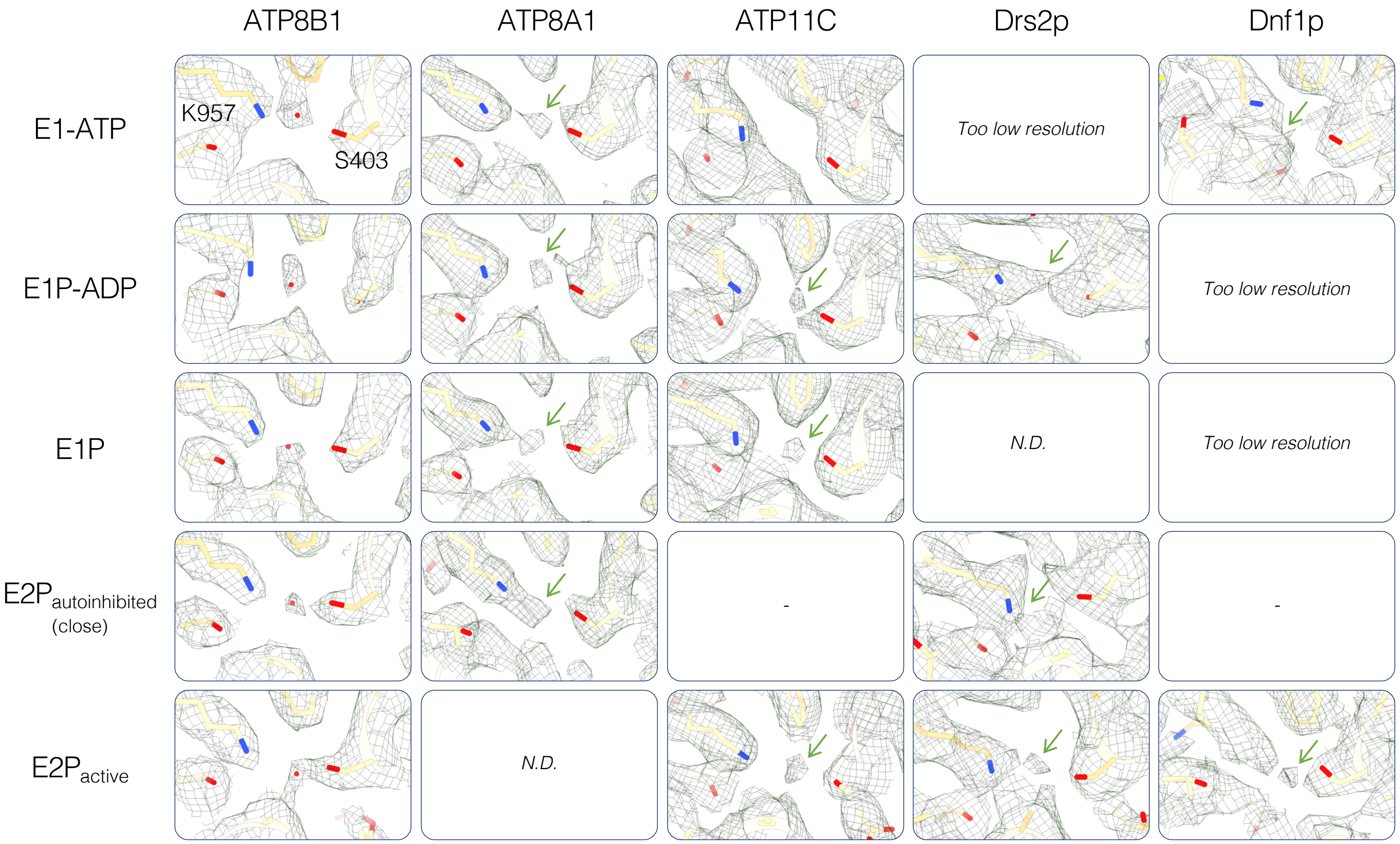


**Figure S4 – Water molecule EM density in the transport site of P4-ATPase**

Comparison of the cryo-EM electron densities of P4-ATPase in E1-ATP, E1P-ADP, E1P and E2P states. For some P4-ATPase the resolution was too low to compared the geometry of the side chains ("too low resolution"), for some the structure has not been determined (N.D.) or do not exist (-). PDB codes: ATP8A1 (6K7J, 6K7K, 6K7N, 6K7L) / ATP11C (7BSP, 7BSQ, 7BSS, 7BSU) / Drs2p (7OH5, 6ROH, 6ROI) / Dnf1p (7WHW, 7DRX).

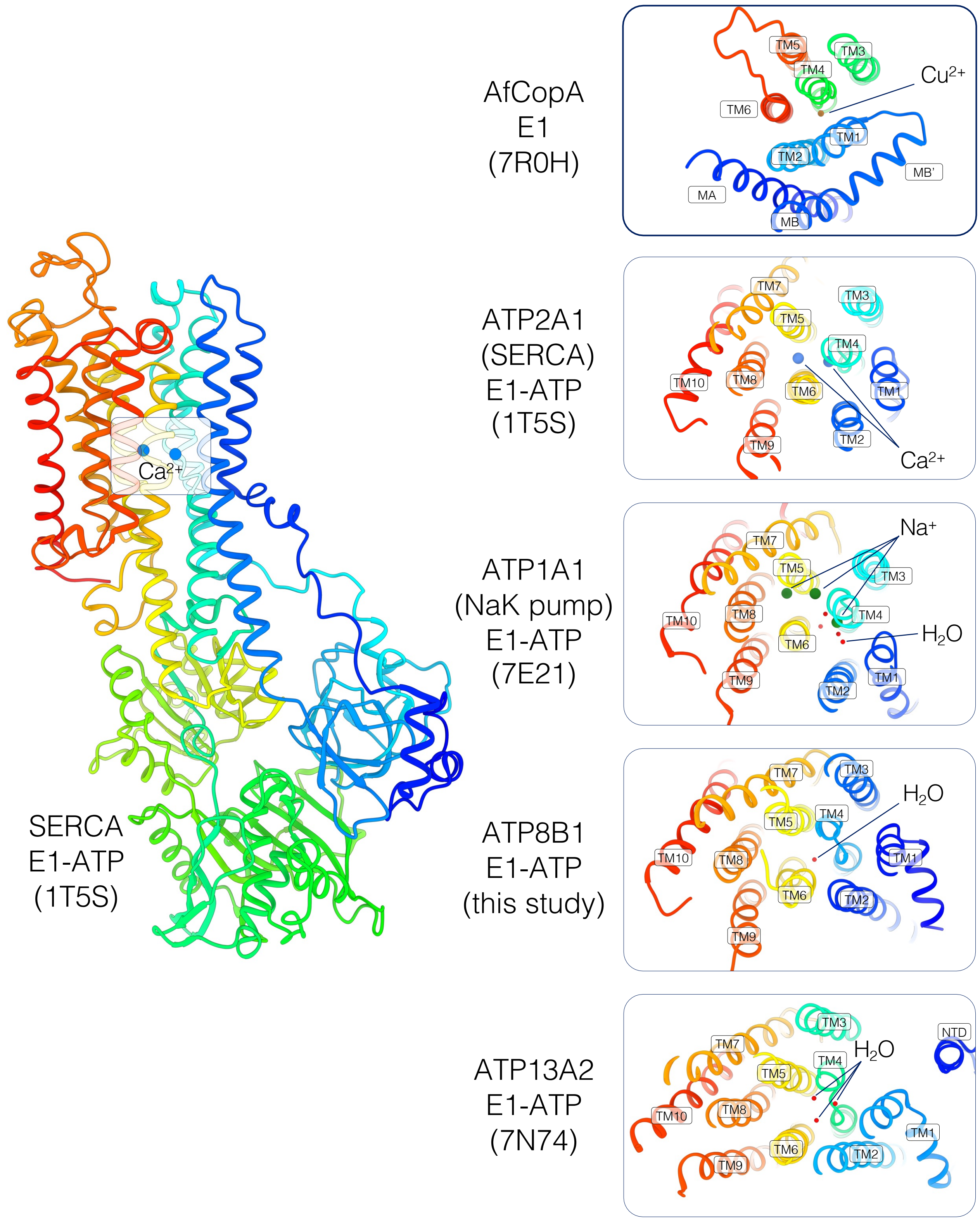


**Figure S5 – Comparison of the canonical transport site in different P-type ATPases in E1 or E1-ATP states.**

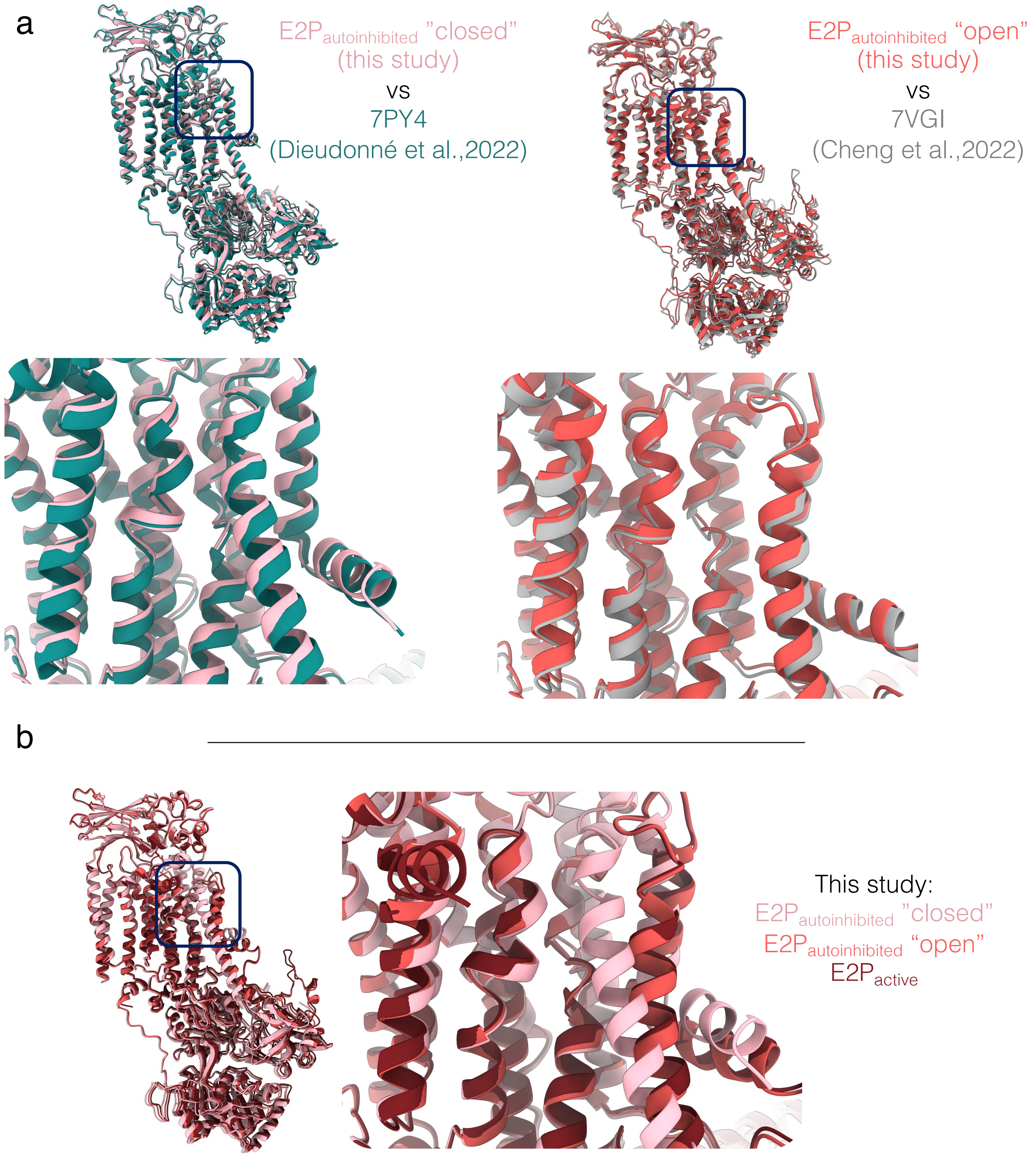


**Figure S6 – Comparison with the previously published structures of ATP8B1-CDC50A autoinhibited states.**

**a)** ATP8B1-CDC50A in the $E2P_{autoinhibited}$ ”closed” conformation (left) is similar to the previously published structure by Dieudonné et al., 2022. ATP8B1-CDC50A in the $E2P_{autoinhibited}$ ”open” conformation (right) is similar to the previously published structure by Cheng et al., 2022. **b)** Structural alignment of the three E2P conformations described in this study. $E2P_{autoinhibited}$ ”open” and the E2P active show a similar TM1-2 open conformation.

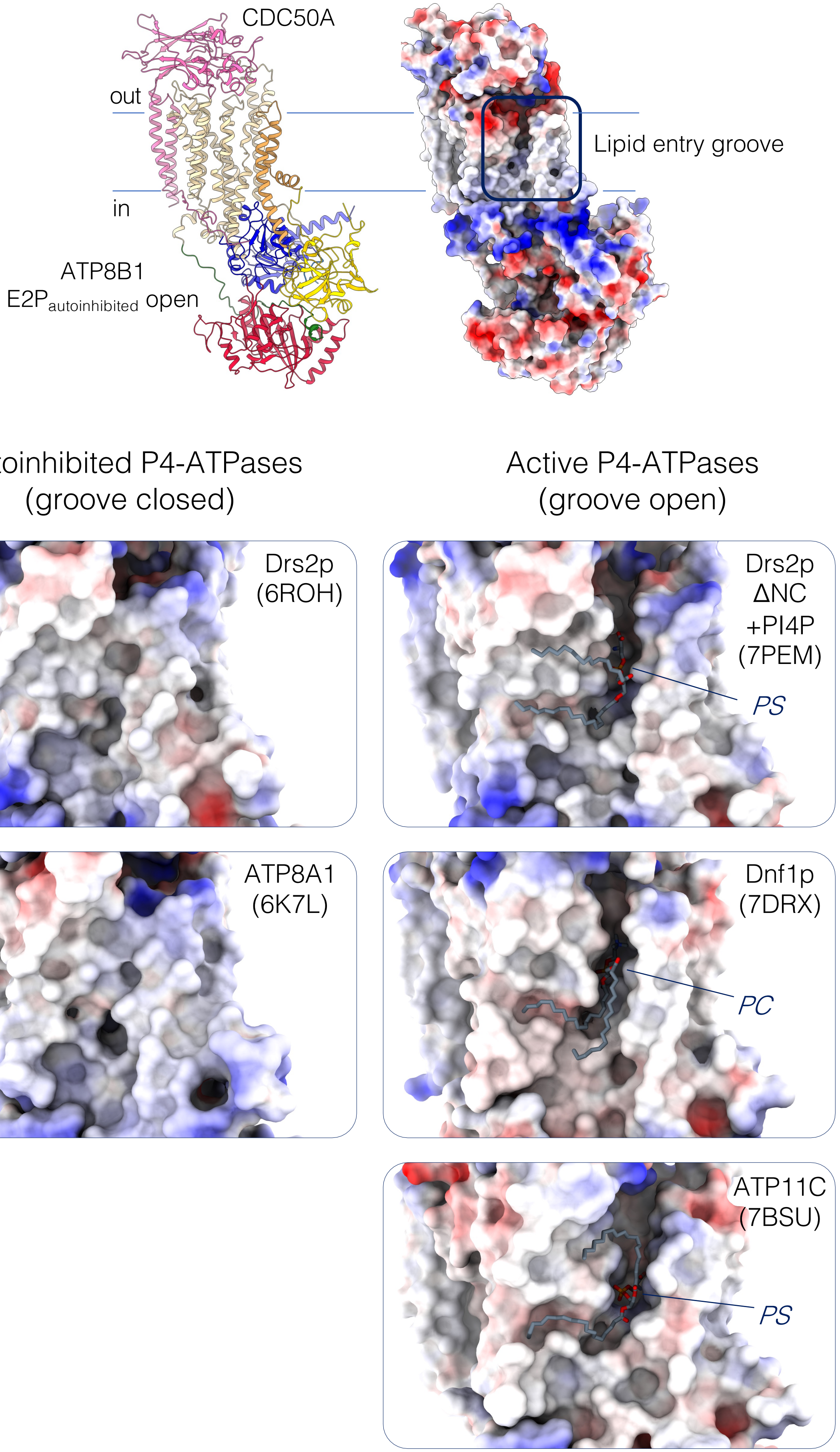


**Figure S7 – Comparison of the lipid groove of autoinhibited and active P4-ATPases in E2P state.**

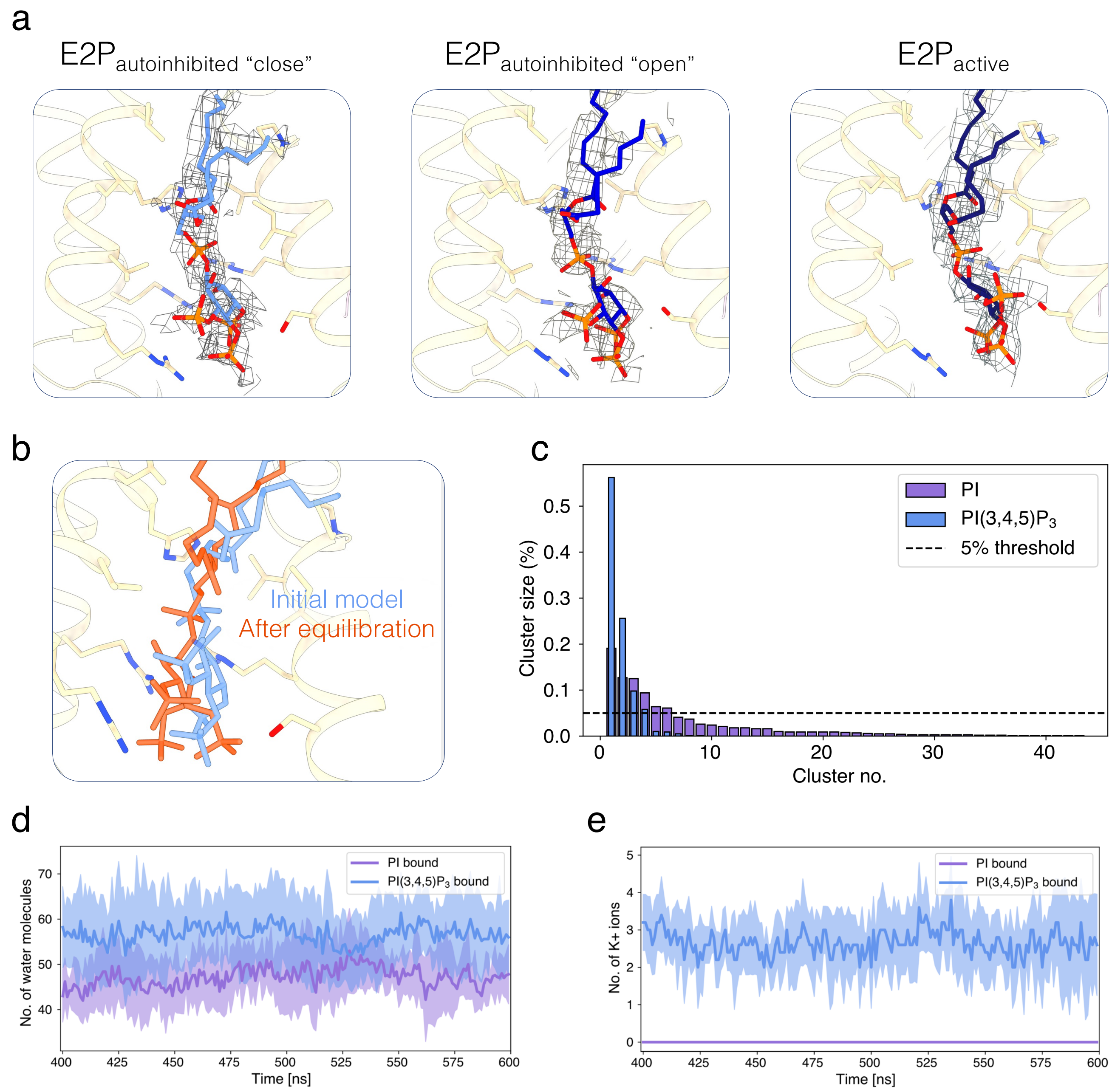


**Figure S8 – PI(3,4,5)$P_3$ binding site in other E2P conformations and complementary analysis of the MD data.**

**a)** PI(3,4,5)$P_3$ binding site and the lipid associated EM density in the three E2P states of ATP8B1 **b)** Comparison of PI(3,4,5)$P_3$ position before (blue) and after (orange) the initial MD equilibration step **c)** All Clusters of inositol ring coordinates from the last 200 ns of each trajectory with Daura's algorithm for PI(3,4,5)$P_3$ (blue) and PI (purple). **d)** Hydration of the lipid binding site. The average number of water molecules present in the lipid binding site for the PI(3,4,5)$P_3$-bound system (blue) and the PI-bound (purple) over the last 200 ns of the simulations. The shaded area indicates the standard deviation over the five replicates of each system. **e)** $K^+$ ions surrounding the lipid headgroup. The average number of $K^+$ ions present within 6 Å of the PI(3,4,5)$P_3$ and the PI (purple) headgroup (blue) over the last 200 ns of the simulations. The shaded area indicates the standard deviation over the five replicates of each system.

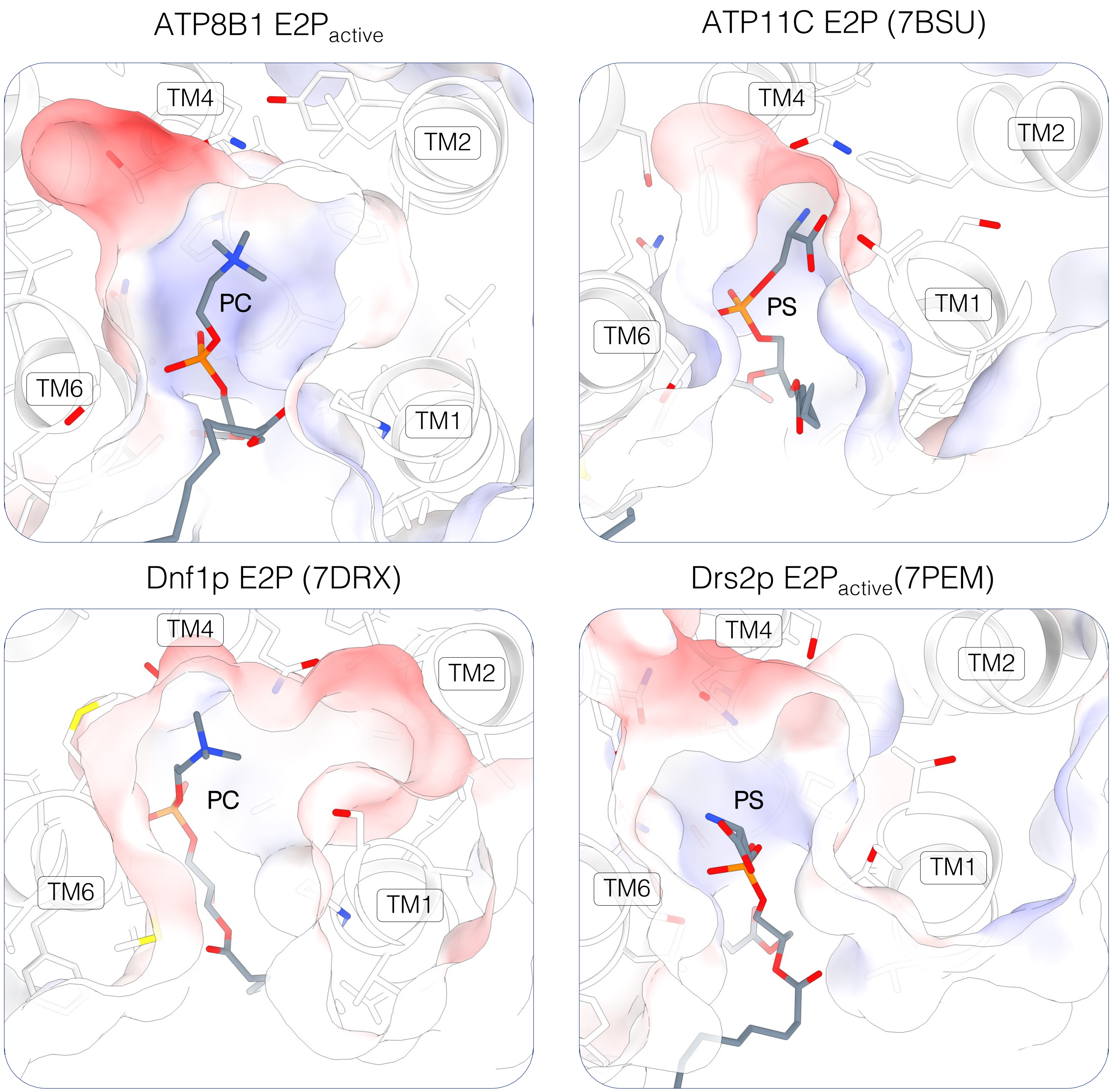


**Figure S9 – Comparison of the lipid binding site in P4-ATPases in E2P conformation**

Exoplasmic view of the transport lipid binding site of different P4-ATPase in E2P conformation. The surface around the transport lipid is shown as electrostatic surface.

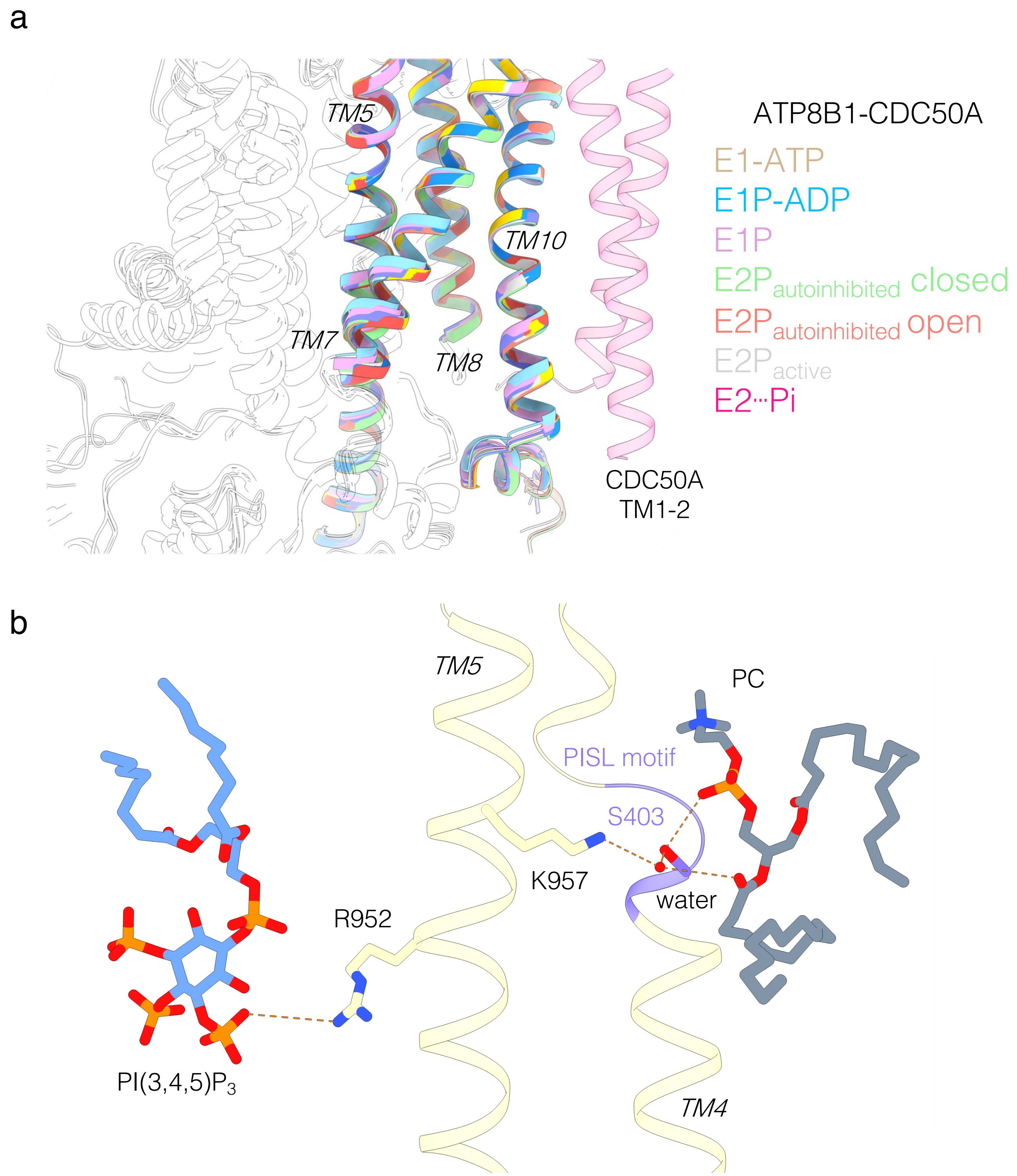


**Figure S10 – PI(3,4,5)$P_3$ binding site in the different ATP8B1-CDC50A conformations reported in this study and its putative link with the lipid transport site. a)** Comparison of the PI(3,4,5)$P_3$ binding site conformation in the different structures presented in this study**. b)** Close-up view of the PI(3,4,5)$P_3$ binding site and the transport lipid binding (PC) site in the E2$P_{active}$ conformation. PI(3,4,5)$P_3$ tightly interacts with R952 on TM5 adjacent to the lipid transport site. For clarity purposes, only TM5 and TM6 are shown.

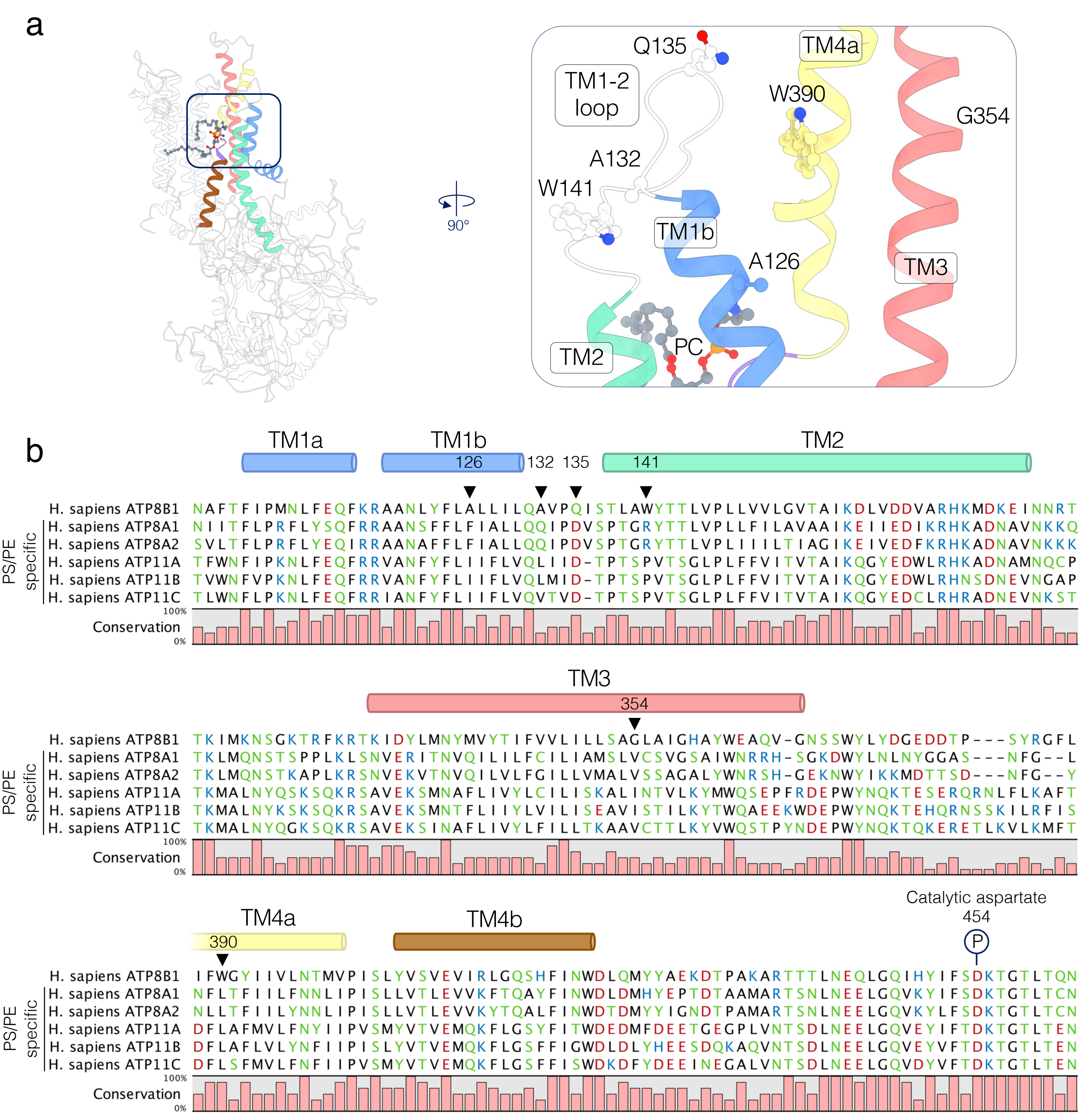


**Figure S11 – Sequence alignment of the TM1-4 of ATP8B1 and PS-specific human P4-ATPases. a)** ATP8B1 in the E2P active PC bound conformation with a close-up view of the TM1-4 organization close to the lipid binding site. TM1, TM2, TM3 and TM4 are blue, turquoise, yellow and red respectively. The non-conserved residues between ATP8B1 and PS-specific human P4-ATPases are shown in sticks. **b)** Sequence alignment of TM1-4 of ATP8B1 and PS-specific human P4-ATPases, colors as in a). (Uniprot ID: hATP8B1(O43520); hATP8A1 (Q9Y2Q0); hATP8A2 (Q9NTI2); hATP11A (P98196); hATP11B (Q9Y2G3); hATP11C (Q8NB49)).

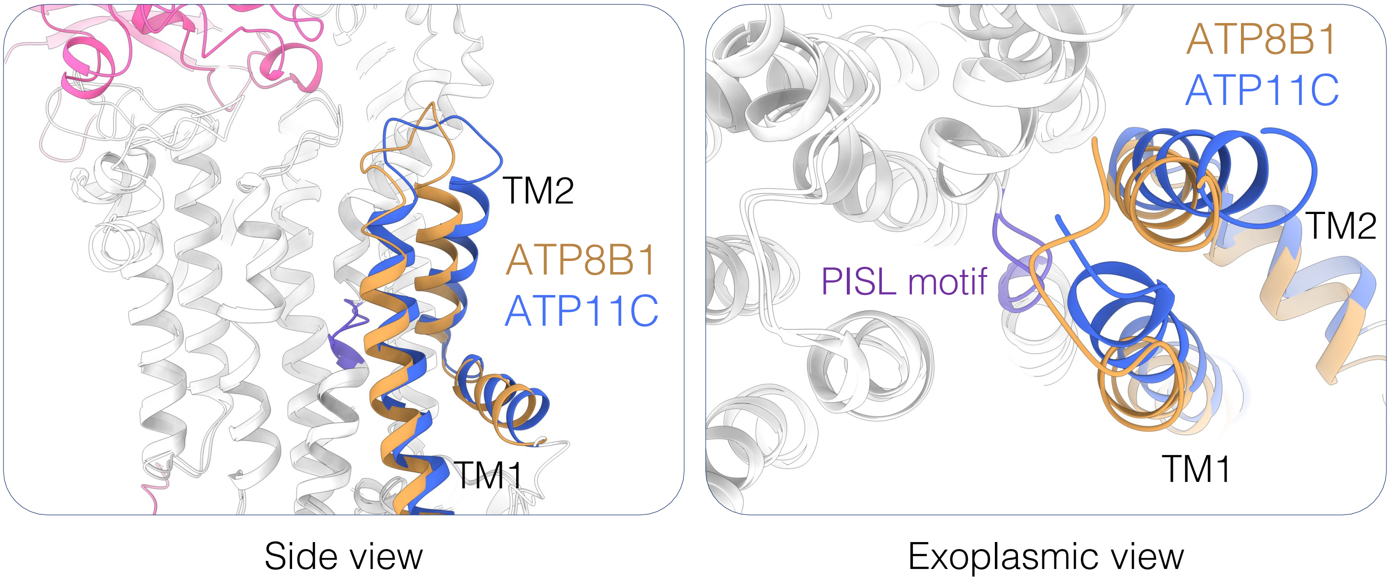


**Figure S12 – TM1-2 orientation in ATP8B1 and ATP11C in E2P substrate bound state**

ATP8B1 TM1-2 are shown in brown, ATP11C (PDB: 7BSU) TM1-2 are shown in blue. The lipid recognition motif (PISL) of TM4 is shown in purple. CDC50A (from the ATP8B1-CDC50A complex) is shown in pink.

Table S1 - Data collection and refinement statistics

| | **E1-ATP** (EMDB-XXXX) (PDB XXXX) | **E1P-ADP** (EMDB-XXXX) (PDB XXXX) | **E1P** (EMDB-XXXX) (PDB XXXX) |
|---|---|---|---|
| **Sample preparation** | | | |
| Construct | ΔCter ATP8B1/CDC50A | ΔCter ATP8B1/CDC50A | ΔCter ATP8B1/CDC50A |
| Ligands / Inhibitors | AMPPCP, PI(4,5)$P_2$ | ADP, $AlF_x$, PI(4,5)$P_2$ | $AlF_x$, PI(4,5)$P_2$ |
| **Data collection and processing** | | | |
| Magnification | | 130,000x | |
| Voltage (kV) | | 300 | |
| Microscope | | Titan Krios G3i | |
| Camera | | Gatan K3 | |
| Physical pixel size (Å/pix) | | 0.647 | |
| Electron exposure (e–/Å$^2$) | | 60 | |
| Defocus range (µm) | | 0.7-1.9 | |
| Number of movies | 7,212 | 10,168 | 10,640 |
| Initial particle images (no.) | 2,486,972 | 3,571,944 | 4,207,324 |
| Final particle images (no.) | 179,570 | 226,859 | 813,354 |
| Symmetry imposed | C1 | C1 | C1 |
| Map resolution (Å) | 3.15 | 2.9 | 2.39 |
| FSC threshold | 0.143 | 0.143 | 0.143 |
| **Refinement** | | | |
| Initial model used (PDB code) | 7PY4 | 7PY4 | 7PY4 |
| Model resolution (Å) | 3.4 | 3.5 | 3.0 |
| FSC threshold | 0.5 | 0.5 | 0.5 |
| Map sharpening *B* factor (Å$^2$) | -100 | -75 | -83 |
| Model composition | | | |
| Non-hydrogen atoms | 9757 | 9755 | 7358 |
| Protein residues | 1198 | 1197 | 903 |
| Water | 1 | 1 | 1 |
| Ligands | 1 ACP, 2 MG, 4 NAG, 1 BMA | 1 AlF, 1 ADP 1 MG, 4 NAG, 1 BMA | 1 AlF, 1 ADP 1 MG, 4 NAG, 1 BMA |
| *B* factors (Å$^2$, min/max/mean) | | | |
| Protein | 5.77/93.41/37.46 | 10.77/90.03/42.76 | 68.52/141.46/92.62 |
| Ligand | 18.68/61.96/43.95 | 33.22/80.69/49.42 | 87.23/122.23/101.94 |
| Water | 13.95/13.95/13.95 | 14.64/14.64/14.64 | 78.48/78.48/78.48 |
| R.m.s. deviations | | | |
| Bond lengths (Å) | 0.003 | 0.003 | 0.003 |
| Bond angles (°) | 0.509 | 0.537 | 0.580 |
| Validation | | | |
| MolProbity score | 1.51 | 1.91 | 1.69 |
| Clashscore | 5.65 | 9.72 | 5.91 |
| Ramachandran plot | | | |
| Favored (%) | 96.79 | 94.01 | 94.73 |
| Allowed (%) | 3.21 | 5.99 | 5.27 |
| Outliers (%) | 0.0 | 0.0 | 0.0 |

| | **E2P$_{autoinhibited}$ (close)** (EMDB-XXXX) (PDB XXXX) | **E2P$_{autoinhibited}$ (open)** (EMDB-XXXX) (PDB XXXX) | **E2P$_{active}$** (EMDB-XXXX) (PDB XXXX) |
|---|---|---|---|
| **Sample preparation** | | | |
| Construct | ATP8B1/CDC50A | ATP8B1/CDC50A | ΔCter ATP8B1/CDC50A |
| Ligands / Inhibitors | ATP, PI(3,4,5)$P_3$ | ATP, PI(3,4,5)$P_3$ | $BeF_x$, PI(3,4,5)$P_3$, POPC |
| **Data collection and processing** | | | |
| Magnification | | 130,000x | |
| Voltage (kV) | | 300 | |
| Microscope | | Titan Krios G3i | |
| Camera | | Gatan K3 | |
| Physical pixel size (Å/pix) | | 0.647 | |
| Electron exposure (e–/Å$^2$) | | 60 | |
| Defocus range (µm) | | 0.7-1.9 | |
| Number of movies | 9,496 | | 9,962 |
| Initial particle images (no.) | 2,982,029 | | 5,471,793 |
| Final particle images (no.) | 573,549 | 51,295 | 256,001 |
| Symmetry imposed | C1 | C1 | C1 |
| Map resolution (Å) | 2.56 | 2.98 | 2.72 |
| FSC threshold | 0.143 | 0.143 | 0.143 |
| **Refinement** | | | |
| Initial model used (PDB code) | 7PY4 | 7PY4 | 7PY4 |
| Model resolution (Å) | 2.8 | 3.2 | 2.9 |
| FSC threshold | 0.5 | 0.5 | 0.5 |
| Map sharpening *B* factor (Å$^2$) | -87 | -66 | -93 |
| Model composition | | | |
| Non-hydrogen atoms | 11757 | 11874 | 11583 |
| Protein residues | 1440 | 1452 | 1411 |
| Water | 1 | 1 | 9 |
| Ligands | 1 IP9,1 MG, 4 NAG, 1 BMA | 1 IP9,1 MG, 4 NAG, 1 BMA | 1 IP9, 1 MG, 1 POV, 1 BEF, 4 NAG, 1 BMA |
| *B* factors (Å$^2$, min/max/mean) | | | |
| Protein | 8.57/88.66/35.12 | 40.84/121.13/70.78 | 28.22/140.18/66.54 |
| Ligand | 24.76/71.63/38.97 | 44.13/107.14/77.92 | 35.48/106.00/62.09 |
| Water | 16.49/16.49/16.49 | 53.89/53.89/53.89 | 34.57/47.42/39.46 |
| R.m.s. deviations | | | |
| Bond lengths (Å) | 0.005 | 0.003 | 0.003 |
| Bond angles (°) | 0.604 | 0.497 | 0.536 |
| Validation | | | |
| MolProbity score | 1.54 | 1.45 | 1.32 |
| Clashscore | 4.78 | 4.40 | 4.67 |
| Ramachandran plot | | | |
| Favored (%) | 95.72 | 96.52 | 97.64 |
| Allowed (%) | 4.28 | 3.48 | 2.36 |
| Outliers (%) | 0.0 | 0.0 | 0.0 |

| | **E2···Pi [PC]** (EMDB-XXXX) (PDB XXXX) | **E2···Pi [PS]** (EMDB-XXXX) (PDB XXXX) | **E2···Pi [PI]** (EMDB-XXXX) (PDB XXXX) |
|---|---|---|---|
| **Sample preparation** | | | |
| Construct | ΔCter ATP8B1/CDC50A | ΔCter ATP8B1/CDC50A | ΔCter ATP8B1/CDC50A |
| Ligands / Inhibitors | $VO_4$, PI(4,5)$P_2$, POPC | $VO_4$, PI(4,5)$P_2$, POPS | $VO_4$, PI(4,5)$P_2$, soy PI |
| **Data collection and processing** | | | |
| Magnification | | 130,000x | |
| Voltage (kV) | | 300 | |
| Microscope | | Titan Krios G3i | |
| Camera | | Gatan K3 | |
| Physical pixel size (Å/pix) | | 0.647 | |
| Electron exposure (e–/Å$^2$) | | 60 | |
| Defocus range (µm) | 0.7-1.9 | 0.7-1.9 | 0.7-1.9 |
| Number of movies | 8,960 | 4,950 (grid #1)<br>6,122 (grid #2) | 10,333 |
| Initial particle images (no.) | 3,120,487 | 4,680,808 | 3,419,101 |
| Final particle images (no.) | 200,083 | 534,468 | 391.193 |
| Symmetry imposed | C1 | C1 | C1 |
| Map resolution (Å) | 2.99 | 2.76 | 2.58 |
| FSC threshold | 0.143 | 0.143 | 0.143 |
| **Refinement** | | | |
| Initial model used (PDB code) | 7PY4 | 7PY4 | 7PY4 |
| Model resolution (Å) | 3.2 | 2.9 | 2.9 |
| FSC threshold | 0.5 | 0.5 | 0.5 |
| Map sharpening *B* factor (Å$^2$) | -85 | -98 | -77 |
| Model composition | | | |
| Non-hydrogen atoms | 11138 | 11139 | 11139 |
| Protein residues | 1363 | 1363 | 1363 |
| Water | 10 | 11 | 11 |
| Ligands | 1 MG, 1 POV, 1 VN4, 4 NAG, 1 BMA | 1 MG, 1 D39, 1 VN4, 4 NAG, 1 BMA | 1 MG, 1 PIE, 1 VN4, 4 NAG, 1 BMA |
| *B* factors (Å$^2$, min/max/mean) | | | |
| Protein | 38.03/200.70/77.71 | 26.94/135.44/58.97 | 39.41/148.73/71.27 |
| Ligand | 43.54/113.55/73.32 | 33.39/93.40/56.30 | 41.91/98.95/69.14 |
| Water | 37.79/63.26/51.22 | 30.62/43.60/36.24 | 44.02/54.97/50.12 |
| R.m.s. deviations | | | |
| Bond lengths (Å) | 0.003 | 0.003 | 0.003 |
| Bond angles (°) | 0.518 | 0.513 | 0.516 |
| Validation | | | |
| MolProbity score | 1.59 | 1.51 | 1.60 |
| Clashscore | 6.07 | 4.72 | 5.94 |
| Ramachandran plot | | | |
| Favored (%) | 96.22 | 96.15 | 96.0 |
| Allowed (%) | 3.78 | 3.85 | 4.0 |
| Outliers (%) | 0.0 | 0.0 | 0.0 |
